# Tumors induce *de novo* steroid biosynthesis in T cells to evade immunity

**DOI:** 10.1101/471359

**Authors:** Bidesh Mahata, Jhuma Pramanik, Louise van der Weyden, Krzysztof Polanski, Gozde Kar, Angela Riedel, Xi Chen, Nuno A. Fonseca, Kousik Kundu, Lia S. Campos, Edward Ryder, Graham Duddy, Izabela Walczak, Klaus Okkenhaug, David J. Adams, Jacqueline D. Shields, Sarah A. Teichmann

**Affiliations:** Department of Pathology, University of Cambridge, Cambridge, CB2 1QP, UK; Wellcome Sanger Institute, Wellcome Genome Campus, Hinxton, Cambridge, CB10 1SA, UK; EMBL-European Bioinformatics Institute, Wellcome Genome Campus, Hinxton, Cambridge, CB10 1SD, UK; Medical Research Council Cancer Unit, Hutchison/Medical Research Council Research Centre, Cambridge, UK; Department of Biology, Southern University of Science and Technology, Shenzhen, China; Iontas Ltd, Iconix Park, Cambridge, CB22 3EG, UK; Theory of Condensed Matter, Cavendish Laboratory, 19 JJ Thomson Ave, Cambridge, CB3 0HE, UK

## Abstract

Tumors subvert immune cell function to evade immune responses, yet the complex mechanisms driving immune evasion remain poorly understood. Here we show that tumors induce *de novo* steroidogenesis in T lymphocytes to evade anti-tumor immunity. Using a novel transgenic steroidogenesis-reporter mouse line we identify and characterize *de novo* steroidogenic immune cells. Genetic ablation of T cell steroidogenesis restricts primary tumor growth and metastatic dissemination in mouse models. Steroidogenic T cells dysregulate anti-tumor immunity, and inhibition of the steroidogenesis pathway was sufficient to restore anti-tumor immunity. This study demonstrates T cell *de novo* steroidogenesis as a mechanism of anti-tumor immunosuppression and a potential druggable target.

## INTRODUCTION

Steroidogenesis is a metabolic process by which cholesterol is converted to steroids^1^. The biosynthesis of steroids starting from cholesterol is often termed “*de novo* steroidogenesis”^1^. Cytoplasmic cholesterol is transported into the mitochondria, where the rate-limiting enzyme CYP11A1 (also known as P450 side chain cleavage enzyme) converts cholesterol to pregnenolone. Pregnenolone is the first bioactive steroid of the pathway, and the precursor of all other steroids (Figure 1a) ^1,2^. The steroidogenesis pathway has been extensively studied in adrenal gland, gonads and placenta. *De novo* steroidogenesis by other tissues, known as “extra-glandular steroidogenesis”, in brain^1,3,4^, skin^5^, thymus^6^, and adipose tissues^7^ has also been reported. Steroid production as a result of immune response in the mucosal tissues, such as in the lung and intestine, has been shown to play a tolerogenic role to maintain tissue homeostasis^8,9^. However the physiological and pathological role of extra-glandular steroidogenesis remains largely unknown^2^.

**Figure 1.**
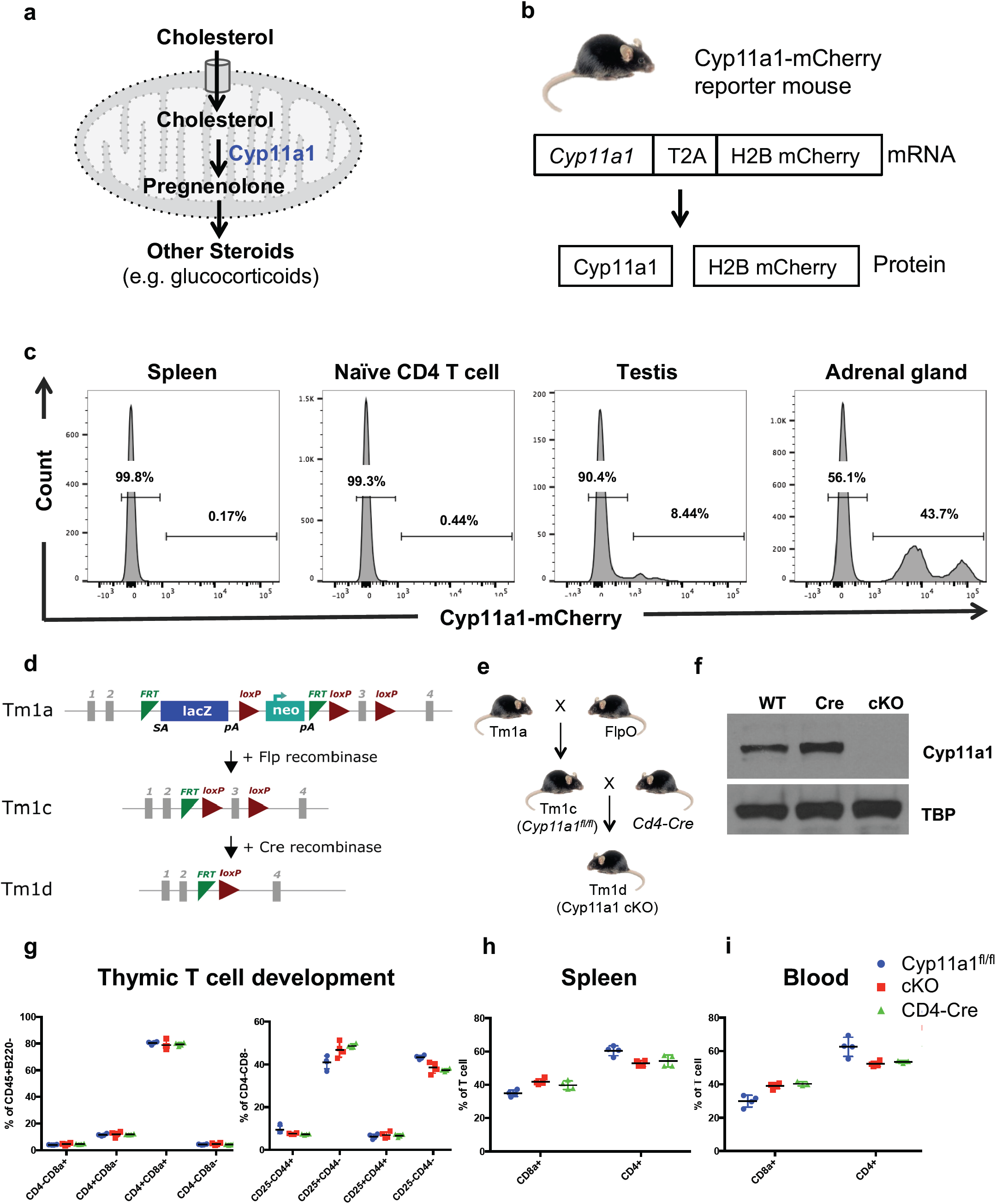
Generation of a de novo steroidogenesis reporter and conditional knockout mice. **a**. Schematic of de novo steroidogenesis pathway. In steroidogenic cells, such as testicular Leydig cells, cholesterol is imported into the mitochondria, where CYP11A1 converts it into pregnenolone, the first bioactive steroid of the pathway and precursor of all other steroids. CYP11A1 is the first and a key rate-limiting enzyme of the pathway. **b**. Cyp11a1-mCherry reporter mice express H2B-tagged mCherry fluorescent protein, and Cyp11a1 as a single mRNA, driven by the endogenous Cyp11a1 promoter. The newly synthesized fusion protein self-cleaves due to presence of a T2A peptide and dissociates into two separate proteins: Cyp11a1 and H2B-mCherry. **c**. Cyp11a1-mCherry reporter mice report Cyp11a1 expression accurately. Spleen, testis and adrenal glands were harvested from Cyp11a1-mCherry reporter mice. Tissues were mechanically dissociated and enzymatically digested into single cell suspension. Splenic naïve CD4^+^ T cells were purified by using anti-CD4 antibody conjugated microbeads by using magnetic activated cell sorting (MACS). Single cell suspensions were analyzed by flow cytometry. Gating: All cells > Singlets > Live cells > Cyp11a1-mCherry. Representative of three independent experiments; each experiment contains 3-4 mice. **d-e**. Generation of a Cyp11a1 conditional knockout mice. Schematic presentation of the Cyp11a1 conditional knockout mouse line and generation of T cell specific Cyp11a1 knockout mice (*Cd4*-Cre;*Cyp11a1*^fl/fl^). The targeting allele is shown in (d) and the crossing strategy is on the (e). **f**. Cyp11a1 knockout efficiency of Cre recombinase in T cells. Splenic naïve T helper cells from cKO (*Cd4*-Cre;Cyp11a1^fl/fl^) mice or control mice (wild type and *Cd4*-Cre) were purified by negative selection using magnetic cell sorting, activated under Th2 differentiation condition, and analysed for Cyp11a1 protein expression by western blot. **g**. Normal thymic development of T cells in Cyp11a1 cKO. Thymus was harvested from Cyp11a1 cKO and control (*Cd4*-Cre and Cyp11a1^fl/fl^) mice, dissociated into single cell suspension, stained with fluorescent conjugated anti-CD4, CD8, B220, CD45, CD25 and CD44 antibodies, and analysed by flow cytometry. Gating: All cells > Singlets > Live cells >CD45^+^ B220^−^ > CD4, CD8 (left panel). CD4^+^CD8^+^ cells represent double positive (DP) stage, CD4^+^CD8^−^ and CD4^−^CD8^+^ cells represent single positive (SP) stage, CD4^−^CD8^−^ cells represent double negative (DN) stage. DN cells of the left panel were gated to show CD25 and CD44 expression to identify DN1 (CD25^−^CD44^+^), DN2 (CD25^+^CD44^+^), DN3 (CD25^+^CD44^−^) and DN4 (CD25^−^ CD44^−^). **h-i**. Normal distribution of Cyp11a1 cKO T cells in the peripheral tissue. Blood and spleen were harvested from Cyp11a1 cKO and control (Cd4-Cre and Cyp11a1^fl/fl^) mice, dissociated into single cell suspension, stained with fluorescent conjugated anti-CD4 and CD8 antibodies, and analysed by flow cytometry.

In cancer, the immunosuppressive tumor microenviroenmnt (TME) prevents immune cells from mounting an effective anti-tumor immune response^10^. Several mechanisms by which cancer cells evade the immune system have been described^11,12^. These include: (1) immune suppression at the TME mediated by immunosuppressive cells, (2) induction of apoptosis in cytotoxic T lymphocytes (CTLs) by the expression of pro-apoptotic ligands e.g., Fas ligand and TRAIL, (3) dysregulating antigen presentation, (4) release of immunosuppressive factors such as IL-10 and TGFβ, and (5) inducing tolerance and immune deviation by mechanisms including, among others, shifting the balance of Th1 immune responses (type 1 immune response) to Th2 (type 2 immune response), and expression of immune inhibitory molecules such as PD-1 (programmed death-1) and CTLA-4 (CTL antigen-4). Established tumor often show type 2 immune responses that drive a immunocompromised and pro-tumorigenic program^13-16^. Further understanding of the anti-tumor immunosuppression would allow us developing new immunotherapies.

Steroid hormones are known immunosuppressive biomolecules^17,18^. We recently reported that type 2 CD4^+^ T cells (Th2 lymphocytes) induce *de novo* steroidogenesis to restore immune homeostasis by limiting the immune response against a worm parasite^19^. Thus we sought to determine whether type 2 T cell-mediated steroidogenesis contributes to the generation of a suppressive niche in the TME.

To study *de novo* steroidogenesis we generated two novel transgenic mouse lines: a fluorescent reporter mouse line (Cyp11a1-mCherry) and a conditional Cyp11a1 (floxed) knockout mouse line. We show the presence of *de novo* steroidogenesis by tumor infiltrating T lymphocytes, but not in unchallenged animals or draining lymph nodes. Genetic ablation of T cell steroidogenesis in Cyp11a1 conditional knockout (cKO) (i.e. *Cyp11a1*^fl/fl^; *Cd4*-Cre) mice restricts experimental primary tumor growth and lung metastasis. We found that intratumoral T cell steroidogenesis dysregulate anti-tumor immunity that could be restored by inhibiting the steroidogenesis pathway pharmacologically. This study therefore demonstrates that T cell *de novo* setroidogenesis is a cause of anti-tumor immunosuppression and a potential drug target for cancer immunotherapy.

## RESULTS

### Generation of Cyp11a1-mCherry reporter and Cyp11a1 conditional knockout mouse line to study *de novo* steroidogenesis

Cyp11a1 expression is known to be a faithful biomarker of *de novo* steroidogenesis due to its role as a first and rate-limiting enzyme in this pathway^1^. Therefore, we generated a novel reporter mouse line to identify Cyp11a1-expressing steroidogenic cells definitively (Figure 1b, c, Extended Data Figure 1a, b, c and d). As expected, mCherry expression was detected in single cell suspensions of testis and adrenal glands but negligible to no expression in the spleen (Figure 1c) or other tissues including lung, kidney, blood, liver, bone marrow, lymph nodes and thymocytes (Extended Data Figure 1b). However, Cyp11a1-mCherry signal was detected specifically in activated type-2 CD4^+^ T helper cells (Th2 cells) upon activation *in vitro* (Extended Data Figure 1c), as reported previously^19^. Cyp11a1 expression was detectable only in mCherry-expressing T helper cells (Extended Data Figure 1d). To date all Cyp11a1-expressing cells have been found to produce at least one steroid^1^. Therefore, we denote Cyp11a1-expressing cells as “*de novo* steroidogenic” or “steroidogenic” cells.

To determine the functional consequences of cell type-specific steroidogenesis we created a *Cyp11a1* floxed (*Cyp11a1*^*fl/fl*^) mouse following EUCOMM/WSI conditional gene targeting strategy^20^. Briefly, a “Knockout-first” (*tm1a*) mouse line was created using a promoter-driven targeting cassette (Figure 1d). The *tm1a* mouse was then crossed with Flp-deleter mice (FlpO) ^21^ to remove the *LacZ* and *Neo* cassette, and generate a *tm1c* allele (i.e. *Cyp11a1*^*fl/fl*^). When crossed with a Cre-driver, the Cre-recombinase removes exon 3 of *Cyp11a1* gene and creates a frameshift mutation (Figure 1d). Because we had initially detected Cyp11a1 expression by Th2 cells^19^, we crossed the *Cyp11a1*^*fl/fl*^ line with a *Cd4*-driven Cre-recombinase to delete *Cyp11a1* and prevent *de novo* steroidogenesis in all T cells (Figure 1e). Deletion efficiency of Cre-recombinase in the *Cyp11a1* cKO (*Cd4-Cre;Cyp11a1*^*fl/fl*^) mice was nearly complete in Th2 cells (Figure 1f). *Cyp11a1* cKO mice showed normal thymic development of T cells, and a normal distribution in the peripheral tissues (Figure 1g, h, i).

### Tumors induce functional Cyp11a1 expression in T cells *in vivo*

Tumor infiltrating T cells are key fate determinants within a tumor, but are often suppressed^22^. The steroidogenesis-inducing type 2 cytokines such as IL4 are also often present in the TME^23,24^, thus we next sought to examine the steroidogenic capacity (i.e. Cyp11a1 induction) of T cells infiltrating tumors, and their impact on tumor development. To explore Cyp11a1 expression *in vivo*, we utilized the well-established B16-F10 melanoma model^25-27^ and implanted tumors subcutaneously in *Cyp11a1*-mCherry reporter mice.

Cyp11a1 expression was detected in immune cells of established primary tumors, but not in tumor-draining brachial lymph nodes (LN) or blood (Figure 2a), indicating that stimulation occurs *in situ*. In support of this, stimulation of splenic and LN resident T cells did not induce Cyp11a1 expression *ex vivo* (Figure 2b, Extended Data figure 2a, LN data not shown). The dominant Cyp11a1^+^ tumor infiltrating immune cells were identified as T cells, predominantly CD4^+^ (helper T cells, Figure 2a). Only a small proportion of the Cyp11a1^+^ cells were reactive to ova, the model antigen in Ova-expressing B16-F10 cells, as determine by staining with MHC-tetramers (Extended Data Figure 2b).

**Figure 2.**
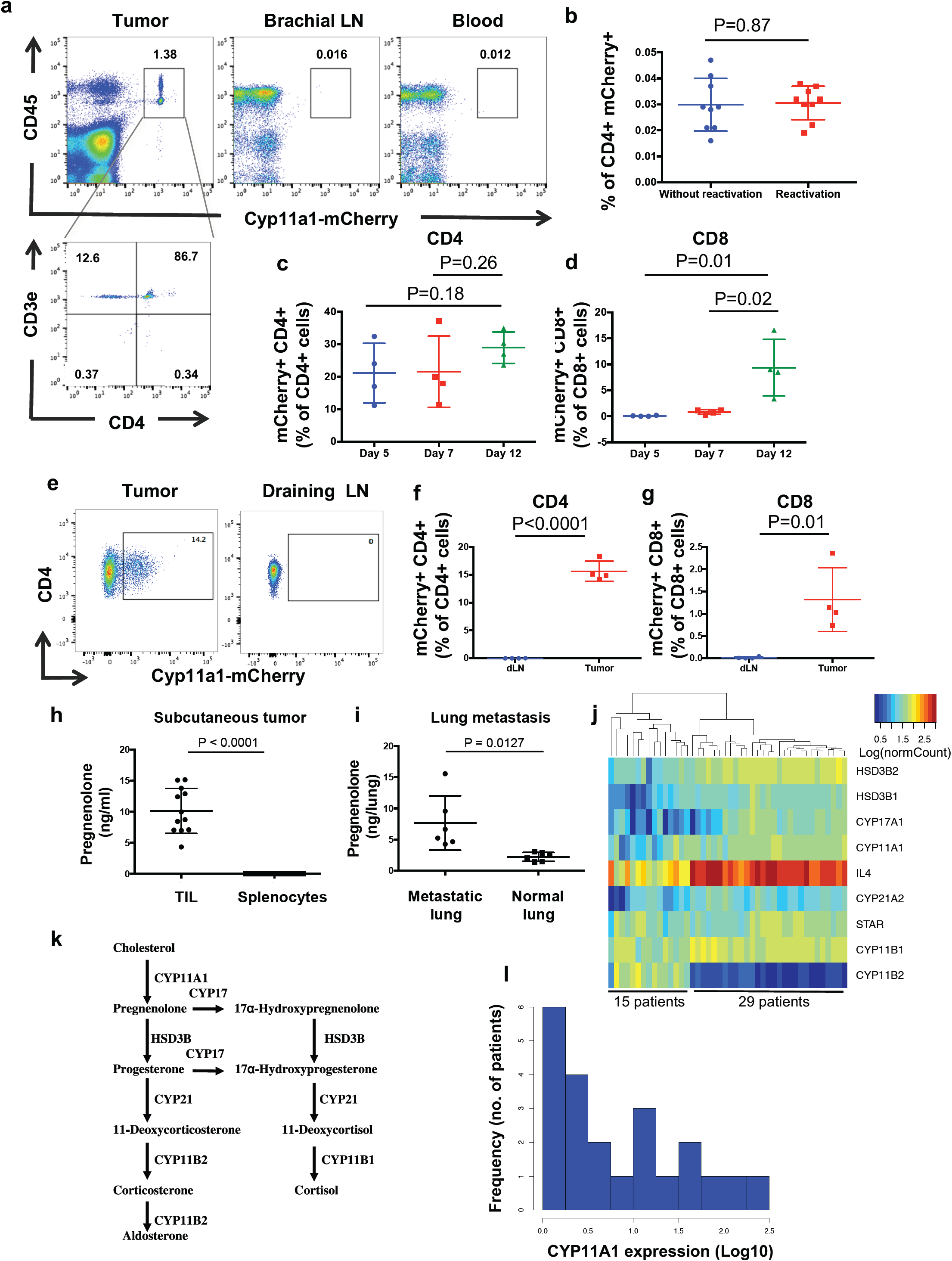
Tumors induce Cyp11a1 expression in T cells in vivo. **a**. Presence of steroidogenic (Cyp11a1^+^) T cells in the B16-F10 melanoma tumor. B16-F10 cells were injected subcutaneously into the shoulder region of Cyp11a1-mCherry reporter mice. After 12 days brachial lymph node (LN), blood and tumor tissues were dissociated into single cell suspensions, and analyzed by flow cytometry. Gating strategy: Singlets > Live cells > CD45, Cyp11a1-mCherry. (N=5) Cyp11a1 expression in CD4^+^ T cells. CD45^+^Cyp11a1-mCherry^+^ cells were further gated to show T helper cell (CD4^+^CD3e^+^) expression of Cyp11a1. **b**. In vitro restimulation of splenic T cells of tumor bearing mice does not induce Cyp11a1 expression. Splenic cells were purified from tumor bearing Cyp11a1-mCherry reporter mice, and reactivated/restimulated in vitro using PMA and ionomycin. Cyp11a1-mCherry expression in CD4^+^ T cells was determined by flow cytometry. P-value was calculated using unpaired two-tailed t-test. Error bars represent mean ± s.d.. **c-d**. *In vivo* intratumoral Cyp11a1 expression dynamics in immune cells. B16-F10 cells were injected subcutaneously into the shoulder region of Cyp11a1-mCherry reporter mice. After 5, 7 and 12 days tumor tissues were dissociated into single cell suspensions, and analyzed by flow cytometry to detect the CD4^+^ T cells (CD45^+^Lin^−^CD4^+^CD3e^+^CD8^−^) (**c**), CD8^+^ T cells (CD45^+^Lin-CD8a^+^CD3e^+^CD4^−^) (**d**) **e-g**. Immune cell-mediated Cyp11a1 expression is conserved in EO771 orthotopic model of triple negative breast cancer. EO771 cells were injected into the mammary fat pad of Cyp11a1-mCherry reporter mice. After 15 days, tumor tissues and tumor draining LN were dissociated into single cell suspensions, and analyzed by flow cytometry to detect the Cyp11a1-mCherry expression in CD4^+^ T cells (CD45^+^Lin^−^CD4^+^CD3e^+^CD8^−^) (**e**,**f**), CD8^+^ T cells (CD45^+^Lin-CD8a^+^CD3e^+^CD4^−^) (**g**) **e**. Representative FACS profile of Cyp11a1^+^CD4^+^ T cells. **f-g**. Representative graphical presentation of one experiment showing Cyp11a1 expressing CD4 and CD8 T cells. **h-i. Tumors induce steroidogenesis in T cells *in vivo*** TIL supernatant contains pregnenolone. B16-F10 tumor-infiltrating leukocytes (TIL) were purified from tumor-bearing mice on post-inoculation day 12, cultured for 48 hours, and the supernatant was analyzed by ELISA to measure pregnenolone. Splenic leukocytes from the tumor bearing mice were used as control. N=12, pooled analysis of three independent experiments, symbol represents individual mouse, error bars represent mean ± s.e.m., unpaired two-tailed t-test. **i**. Metastatic dissemination of cancer cells induces steroid biosynthesis. Metastasized lungs were harvested 10 days post-B16-F10 intravenous injection in C57BL/6 mice, dissociated into single cell suspension, cultured for 48 hours and the supernatant was analyzed by ELISA. Naïve uninfected lungs (normal lung) were used as control. N=6, symbol represents individual mouse, pooled analysis of two independent experiments, error bars represent mean ± s.e.m., unpaired two-tailed t-test. **j**. Human tumors induce de novo steroidogenesis. Steroidogenic gene expression is correlated with IL4 expression in human melanoma. Hierarchical clustering of steroidogenic genes and IL4 mRNA expression across 44 melanoma patient samples (Raw data source: GEO: GSE19234). **k**. Schematics showing glucocorticoids (corticosterone and cortisol) synthesis pathway. **l**. Frequency distribution histogram showing CYP11A1 mRNA expression (normalized read counts, log10 scale) across 22 melanoma patients’ tumor infiltrating CD4^+^ T cells (Raw data accession code EGAD00001000325).

Having identified presence of Cyp11a1-expressing T cells in established tumors, we next examined expression dynamics during tumor progression. Using the Cyp11a1-mCherry reporter mice we observed an expression of Cyp11a1 in CD4^+^ T cells which remained stable through the development of the tumor through days 7 and 12. By contrast, CD8^+^ T cells only expressed Cyp11a1 by day 12 (Figure 2c, d). Expression dynamics of minor and rare Cyp11a1^+^ non-T cells (mast cell and basophil) is shown in Extended Data Figure 2c, d.

To determine whether Cyp11a1 induction in T cells is conserved in other tumor types we analyzed a EO771 orthotopic model of breast cancer^28-30^ (Figure 2e-g, Extended Data Figure 2e). Again, we found that Cyp11a1 was upregulated in tumor infiltrating T cells (Figure 2e, f, g), but not in tumor draining lymph node or spleen. We also detected minor and rare population of other Cyp11a1^+^ type-2 immune cells, such as mast cells and basophil (Extended Data Figure 2e).

Next, we sought to measure the functional output of Cyp11a1 expression. Significant concentrations of the steroid pregnenolone were detected exclusively in immune cells isolated from tumors, with negligible levels detected in cells from the spleen (Figure 2h). Using the B16-F10 model of experimental metastatic dissemination^31^, we determined that lungs with metastatic nodules, but not control lungs without metastatic nodules, had elevated levels of pregnenolone (Figure 2i).

The presence of tumor cells did not induce or augment the Cyp11a1 expression in T cells *in vitro* (Extended Data Figure 2f). This indicates that type-2 (Th2) cytokines within the TME induce Cyp11a1 expression by T cells and this cannot readily be mimicked *in vitro*.

Having observed steroidogenic T cells in murine melanoma, we turned to publicly available transcriptomic data sets to verify our findings and ascertain relevance in the human setting. We identified *CYP11A1* mRNA expression, and thus steroidogenic potential, in a range of cancer types including liver, breast, prostate, lung, kidney, sarcoma, glioma, uterine, cervical, lymphoma and melanoma (Figure 2j and Extended Data Figure 2g, h)^24^. Human melanoma tissues represented a prominent steroidogenic tumor type, expressing *CYP11A1, HSD3B1, HSD3B2, CYP17A1, CYP21A1, CYP11B1* and not expressing *CYP11B2* (Figure 2k). Together this was indicative of melanoma-driven production of glucocorticoids (Figure 2k, Extended Data Figure 2i).

Interestingly, in melanoma, steroidogenic gene expression was correlated with *IL4* expression (Figure 2k, Extended Data Figure 2j), a key inducer of T cell steroidogenesis^19^. Moreover, analysis of human tumor infiltrating CD4^+^ T cell transcriptomes, confirmed *CYP11A1* expression (Figure 2l) implying that CD4^+^ T cells are a source of steroids in tumors, mirroring the murine setting. Collectively these data indicate that TILs produce steroids within the tumor both in human and mouse.

### Ablation of T cell de novo steroidogenesis restricts experimental tumor growth and metastasis

To determine the functional consequences of T cell-driven steroidogenesis in tumors we subcutaneously implanted *Cyp11a1* cKO mice with B16-F10 cells. Genetic deletion of *Cyp11a1* in T cells significantly restricted primary tumor growth rates and final volumes (Figure 3a). Similarly, in the experimental metastasis model, impaired lung colonization was observed in the absence of T cell-expressed Cyp11a1. There was a significant reduction in number of metastatic foci in lungs of *Cyp11a1* cKO mice compared to the control mice (Figure 3b). In the B16-F10 subcutaneous melanoma model, topical application of exogenous pregnenolone at the primary tumor site was sufficient to compensate for the Cyp11a1 deficiency, restoring tumor growth to levels comparable with control mice (Figure 3c). This suggests that the tumor restriction was a consequence of the absence of pregnenolone synthesis. Similar results were obtained using the EO771 breast cancer orthotopic tumor model: deletion of *Cyp11a1* in T cells restricted dissemination to and nodule formation in the lung (Figure 3d). Importantly, in both tumor models, pharmacological inhibition of Cyp11a1 by aminoglutethimide (AG) was sufficient to recapitulate the tumor restriction phenotype resulting from *Cyp11a1* genetic ablation (Figure 3e, f). Together, these data indicate that, T cell-derived steroids can support tumor growth and both genetic and pharmacological interference with the pathway can restrict tumor growth.

**Figure 3.**
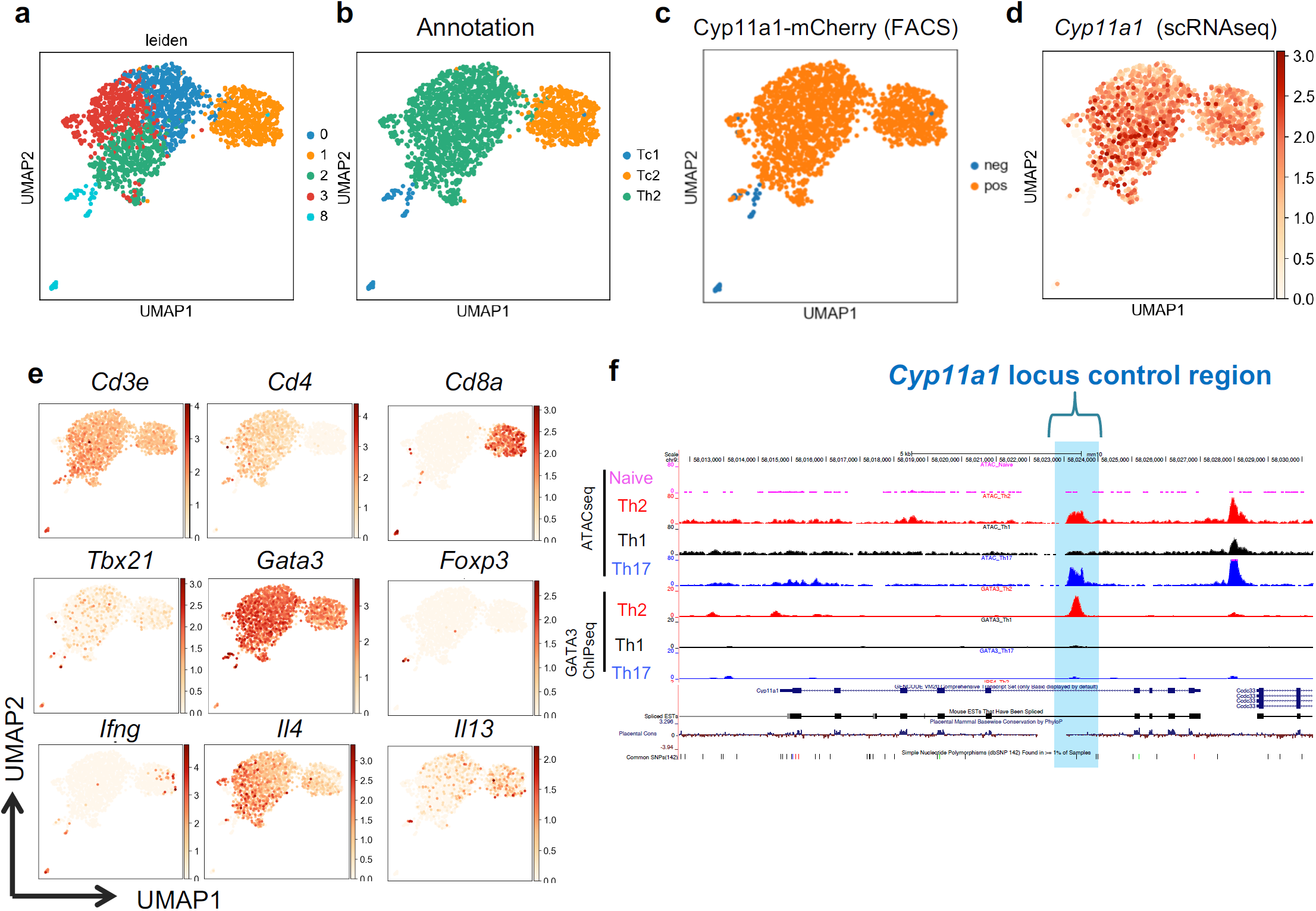
Single-cell transcriptomics revealed gene expression identity and gene expression pattern of intratumoral steroidogenic T cells. **a**,**b**. UMAP visualization of the tumor infiltrating T cells (a) with annotations of the clusters (b). Tc1 represents type-1 CD8^+^ T cells, Tc2 represents type-2 CD8^+^ T cells and Th2 represents type-2 CD4^+^ T cells. **c**,**d**. mCherry protein expression perfectly reports *Cyp11a1* mRNA expression. Cyp11a1-mCherry protein expression according to the FACS data (c). *Cyp11a1* mRNA expression in the scRNA-seq data (d). **e**. Expression of cell type specific signature genes that were used to annotate the clusters. **f**. Determination of *Cyp11a1* locus control region. Open chromatin regions were identified by ATAC-seq of T helper cells. In vitro generated Th2 and Th17 cells were used as positive control. Naïve and Th1 cells were used as negative control. Gata3 ChIP-seq data were analyzed to determine the binding site. The open chromatin region where Gata3 occupy is highlighted blue and considered as locus control region.

### Inhibition of T cell steroidogenesis stimulates anti-tumor immunity

Steroid hormones can induce an immunosuppressive M2-like phenotype in macrophages^17,32,33^ cell death and anergy in T cells^17 34-36^, tolerance in dendritic cells^17,37-39^ and increase the frequency of Treg cells^40-42^. Therefore, we tested whether steroidogenic T cells support tumor growth through the induction of immunosuppressive phenotypes in infiltrating immune cells. To determine whether intratumoral macrophages were M1 or M2 type, we purified tumor infiltrating macrophages and analyzed mRNA expression of the M2-macrophage signature genes *Arg1* and *Tgfb1*. In tumor-infiltrating macrophages from *Cyp11a1* cKO mice, *Arg1* and *Tgfb1* mRNA expression was significantly reduced compared to the control mice, indicating fewer of the tumor-supporting M2 macrophages in the T cell-specific *Cyp11a1* knockout mice (Figure 4a). We confirmed changes in macrophage characteristics at the protein level, showing a significant decrease in Arg1^+^ macrophages (M2) with coincident increase in tumor infiltrating iNOS^+^ macrophages (M1) upon T cell-specific *Cyp11a1* deletion (Figure 4b, c). Hence, depletion of Cyp11a1 in T cells increases the M1/M2 macrophage ratio in the TME.

**Figure 4.**
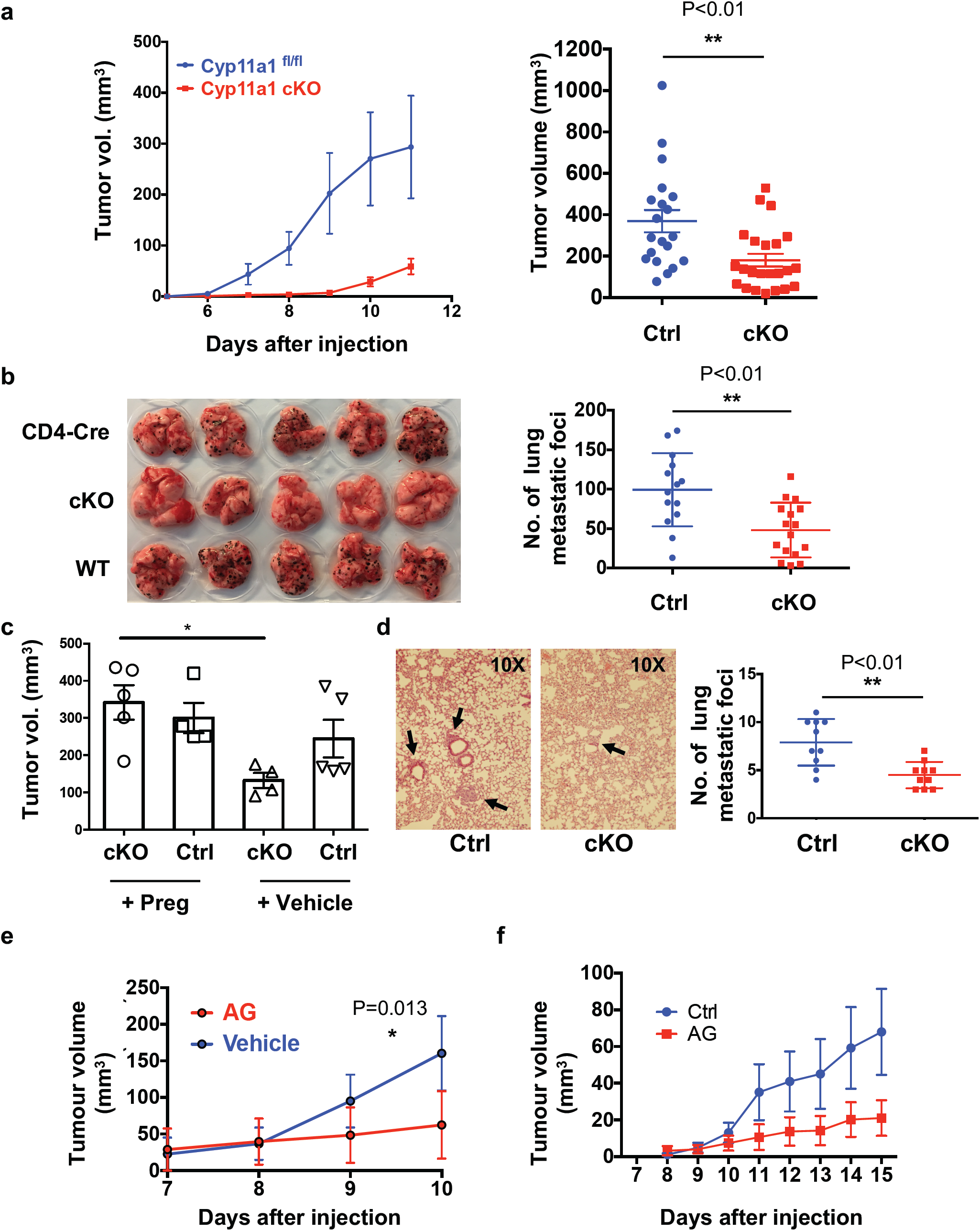
Ablation of T cell de novo steroidogenesis restricts experimental tumor growth and metastasis. **a**. Genetic deletion of Cyp11a1 in T cells inhibits primary tumor growth of B16-F10 melanoma cells. Left-panel: B16-F10 subcutaneous tumor growth curve assessed in T cell specific Cyp11a1 cKO (cKO) and Cd4-Cre control mice (n=5, representative of four independent experiments, error bars represent mean ± s.e.m.). Tumors were measured every day from 6th day after subcutaneous injection of B16-F10 cells. Right panel: Graphical presentation of end-point tumor volume (n ≥ 20, pooled data of four independent experiments, error bars represent mean ± s.e.m., unpaired two-tailed t-test). **b**. Genetic deletion of Cyp11a1 in T cells inhibits experimental lung metastasis of B16-F10 melanoma. Left panel: Representative photograph of pulmonary metastatic foci produced 10 days after tail vein injection of B16-F10 cells. Right panel: Graphical presentation of the numbers of lung metastatic foci (right panel) (n ≥ 15, pooled data of three independent experiments, error bars represent mean ± s.d., unpaired two-tailed t-test). **c**. Pregnenolone complements the Cyp11a1 deficiency in Cyp11a1 cKO mice. Cyp11a1 cKO and Cd4-Cre control mice were injected with B16-F10 cells. Pregnenolone or vehicle (DMSO) applied topically at the primary tumor site every 48 hrs. Tumor volume was measured at the end-point at day 12. N=5, error bars represent mean ± s.e.m., one way ANOVA, representative experiment. **d**. Genetic deletion of *Cyp11a1* in T cells inhibits experimental lung metastasis of EO771 breast cancer cells. Representative hematoxylin and eosin stained histologic photograph of pulmonary metastatic foci produced 10 days after tail vein injection of EO771 cells in Cyp11a1 cKO and Cyp11a1^fl/fl^ mice. Graphical presentation of the numbers of lung metastatic foci (right panel). **e**. Cyp11a1 inhibitor aminoglutethimide restricts melanoma tumor growth. B16-F10 cells were injected subcutaneously in C57BL/6 mice with or without Cyp11a1 inhibitor aminoglutethimide (AG). AG treatment was continued with a 48 hrs interval (N=5, error bars represent mean ± s.e.m, two way ANOVA, P=0.013, representative experiment). **f**. Cyp11a1 inhibitor aminoglutethimide restricts orthotopic breast cancer growth. EO771 cells were injected into the mammary fat pad in C57BL/6 mice with or without Cyp11a1 inhibitor AG. AG treatment was continued with a 48 hrs interval. (N=5, error bars represent mean ± s.e.m., representative experiment).

Consistent with enhanced tumor immunity, we found significantly higher levels of the inflammatory cytokines *Ifng* and *Tnfa* expression were identified in CD8^+^ T cells following *Cyp11a1* ablation (Figure 4d). Moreover the expression of the co-inhibitory receptors^43-45^ PD1 and TIGIT on tumor-infiltrating T cells was reduced in CD4^+^ TILs in the *Cyp11a1* cKO mice compared with control littermates (Extended Data Figure 3a). The data indicate greater T cell functionality and less exhaustion in the *Cyp11a1* cKO mice.

To test cytotoxic capacity of T and NK cell populations, we examined the degranulation response of these cells by analyzing cell surface expression of CD107a/LAMP1 in tumor-infiltrating T and NK cells. We observed a significantly increased proportion of degranulating CD107a^+^CD8^+^ T cells in *Cyp11a1* cKO mice compared to control mice (Figure 4e). The degranulation response in NK cells and CD4^+^ T cells was also enhanced (Figure 4f, g).

Finally, we observed that proportion of intratumoral Tregs was decreased (Figure 4h). Altogether these data suggest that inhibition of T cell steroidogenesis by genetic deletion of *Cyp11a1* changes the immune cell composition in the tumor microenvironment in favour of anti-tumor immunity.

Similar to the genetic deletion of *Cyp11a1*, application of the Cyp11a1 inhibitor aminoglutethimide significantly increased frequencies of M1 macrophages and degranulating T and NK cells (Figure 4i, j, k, l, Extended Data Figure 3b) recapitulating anti-tumor phenotypes. *Cyp11a1* inhibition also decreased the number of M2 macrophages in the tumor (Figure 4i). Together, these data demonstrate the potential to stimulate anti-tumor immunity through pharmacological suppression of *Cyp11a1-*dependent steroidogenesis pathways.

## DISCUSSION

The endocrine importance of systemic steroid hormones is well documented in regulating cell metabolism and immune cell function but the intracrine, autocrine and paracrine role of local cell type specific steroidogenesis is less clear, particularly in pathologies such as cancer^1,2^. This is in part due to the lack of tools to study steroidogenesis in a tissue-specific manner *in vivo*. To overcome this, we generated a novel *Cyp11a1*-mCherry reporter and a conditional *Cyp11a1* knockout mouse strain to identify *de novo* steroidogenic cells and study their role *in vivo*. Using these discovery tools, complemented by pharmacological intervention, we uncovered a novel anti-tumor immune suppression mechanism that may be exploited clinically to boost the anti-tumor immunity (Figure 5).

**Figure 5.**
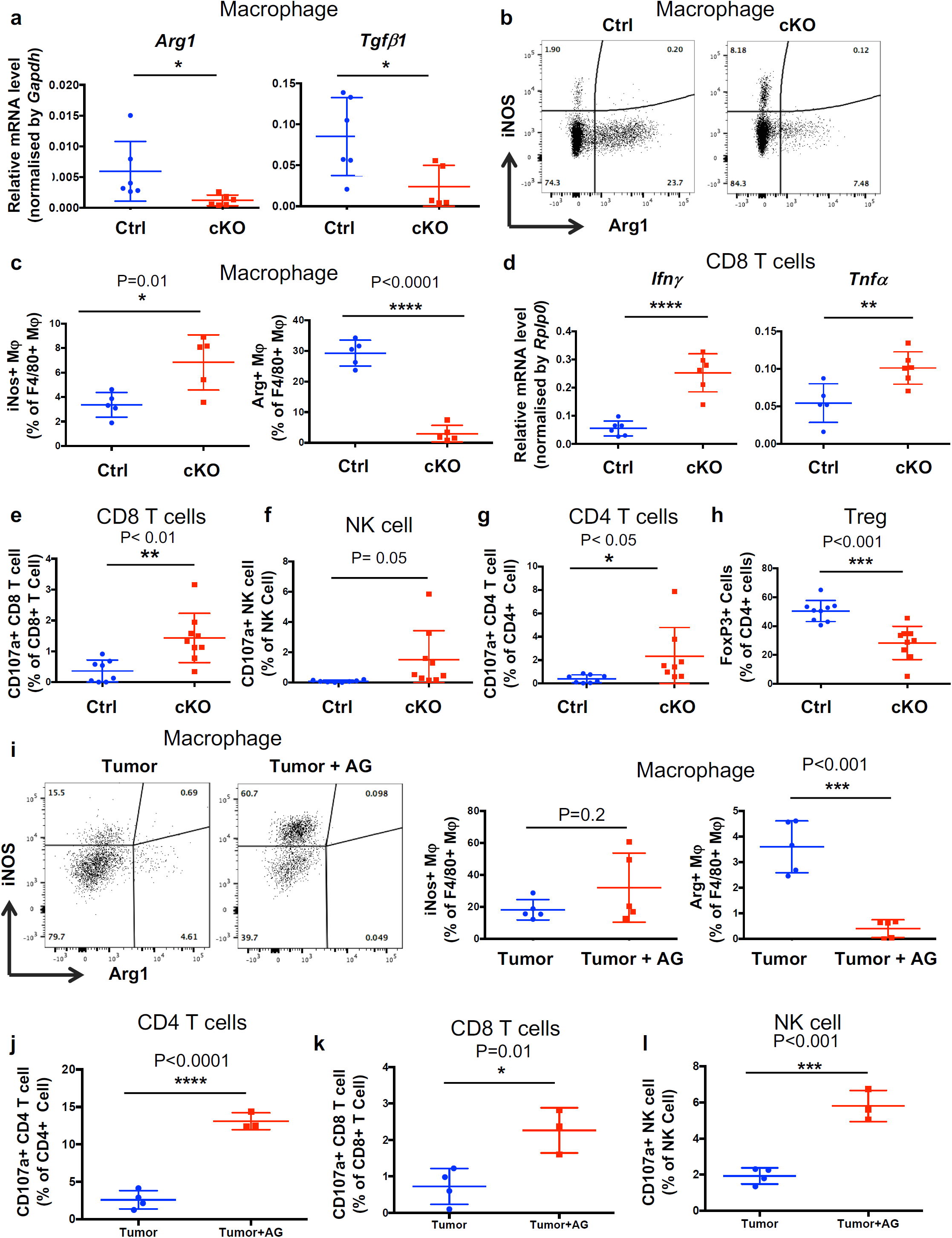
Inhibition of T cell steroidogenesis stimulates anti-tumor immunity. **a-h**. Comparing Cyp11a1 cKO and *Cd4*-Cre control mice with B16-F10 cells injected subcutaneously (n= 5 or 6, error bars represent mean ± s.d., unpaired two-tailed t-test, representative experiment). **a**. Tumor infiltrating macrophages (Lin-CD11b^+^) were purified by cell sorting at day 12. *Arg1* and *Tgfb1* mRNA expression was quantified by RT-qPCR, with mRNA expression level normalized by *Gapdh* mRNA expression. **b-c**. Tumor infiltrating macrophages (Lin^−^CD11b^+^F4/80^+^) analyzed by flow cytometry at day 12 to examine iNOS and Arg1 expression. Representative FACS profile of the expression (b). Representative graphical representation of one experiment (n = 5). **d**. Tumor infiltrating CD3e^+^CD8^+^ T cells purified by cell sorting at day 12, reactivated ex vivo, and *Ifng* and *Tnfa* mRNA expression quantified by RT-qPCR, with mRNA expression level was normalised by *Rplp0* expression. **e**. Cytotoxic T lymphocyte (CD8^+^ T cell) degranulation assay. CD107a/LAMP1 expression on tumor infiltrating CD8^+^ T cells was analyzed by flow cytometry after 12 days post-inoculation of B16-F10 cells. Gating: All cells > singlets > live cells > CD8^+^ T cell > CD107a. **f**. NK cell degranulation assay by measuring CD107a/LAMP1 expression on tumor infiltrating NK cells. Gating: All cells > singlets > live cells > NK cells > CD107a. **g**. CD4^+^ T cell degranulation assay by measuring CD107a/LAMP1 expression on CD4^+^ T cells. Gating: All cells > singlets > live cells > CD4^+^ T cell > CD107a **h**. Number of regulatory T cells in the TME decreases upon Cyp11a1 deletion. Flow cytometric analysis to show the changes in B16-F10 subcutaneous tumor infiltrating Treg (CD4^+^CD3e^+^FoxP3^+^) population upon Cyp11a1 deletion. **i-l**. Comparing immunophenotype in Cyp11a1 inhibitor, AG, treated and untreated mice with B16-F10 cells injected subcutaneously (n= 5 or 6, error bars represent mean ± s.d., unpaired two-tailed t-test, representative experiment)., representative experiment). **i**. Representative FACS profile to show iNOS and Arg1 expression in tumor infiltrating CD45^+^Lin^−^ CD11b^+^F4/80^+^ macrophages (left panel). Graphical presentation the iNOS and Arg1 expression of a representative experiment (right panel). **j**. CD4^+^ T cell degranulation assay by measuring CD107a/LAMP1 expression on CD4^+^ T cells. Gating: All cells > singlets > live cells > CD4^+^ T cell > CD107a. **k**. Cytotoxic T lymphocyte (CD8^+^ T cell) degranulation assay. CD107a/LAMP1 expression on tumor infiltrating CD8^+^ T cells was analyzed by flow cytometry after 12 days post-inoculation of B16-F10 cells. Gating: All cells > singlets > live cells > CD8^+^ T cell > CD107a. **l**. NK cell degranulation assay by measuring CD107a/LAMP1 expression on tumor infiltrating NK cells. Gating: All cells > singlets > live cells > NK cells > CD107a.

**Figure 6.**
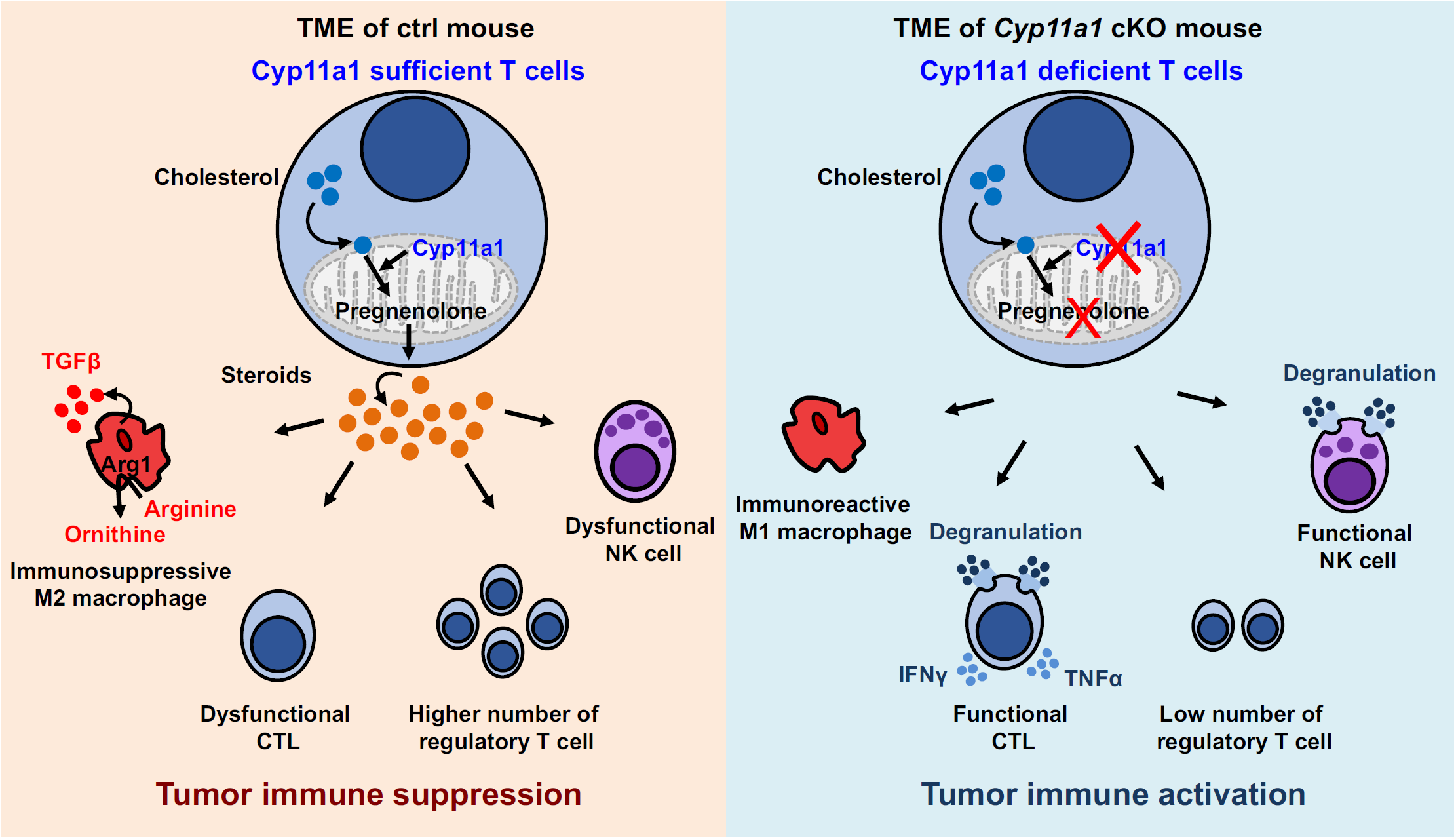
Graphical abstract of the discovery. T cell mediated de novo steroidogenesis in the tumor microenvironment promotes tumor growth by inhibiting anti-tumour immunity, in part by inducing M2 phenotype in macrophages and suppressing T and NK cell function. Genetic deletion of Cyp11a1 or pharmacologic inhibition of Cyp11a1 activity stimulates anti-tumor immunity by increasing the number of M1 macrophages, functional T cells and decreasing the number of regulatory T cells, M2 macrophages.

The TME harbors innate immune cells such as macrophages, neutrophils, eosinophils, mast cells, myeloid derived suppressor cells, dendritic cells, and natural killer cells, adaptive immune cells such as T and B cells, stromal cells such as fibroblasts, endothelial cells, pericytes, mesenchymal cells, and cancer cells. The presence of immune cells in the tumor microenvironment raised a long-standing question: How tumors evade the immune surveillance and anti-tumor immunity? The present understanding suggests that the balance between protective anti-tumoral immune cells (and other protective factors) and pro-tumoral immunosuppressive cells (and other immunosuppressive factors) determines the fate. Anti-tumoral immune cell subsets includes Th1, Tc1 (CD8^+^ CTL), M1 macrophage, DC1 which are antagonized by Th2, M2, tolerogenic DC, immature DC, DC2, Treg and myeloid derived suppressor cells. Established tumors are often foster immunosuppressive and tolerogenic immune cells. In this report, the discovery of immune cell-mediated steroid biosynthesis in the TME explains beautifully how immunosuppressive TME can be achieved. Previously it has been shown in several independent reports that steroids induce anergy and cell death in T cells^34-36,46^, M2 phenotype in macrophages^32,33^, tolerance in DCs^37-39^ and increase the frequency of Treg cells^40-42^. Here we demonstrate that tumor infiltrating T cells can produce steroids and that the inhibition of T cell-mediated steroid biosynthesis tips the balance toward protective immunity. Importantly, this pathway can be targeted pharmacologically. This may be exploited therapeutically in three ways: (1) steroidogenic gene expression or the presence of steroids might be used to stratify the patients (2) use of pharmacologic drug (i.e. anti-steroidogenic) can convert the TME into protective type-1 inflammatory condition by reducing the steroid level and (3) immunotherapy should work better in absence of immunosuppressive steroids.

We identified three distinct classes of immune cells with steroidogenic potential, T cells (Th2 and Tc2, major population), mast cell (rare and minor population), basophil (rare and minor population) and a few monocyte/macrophage. T cells outnumbered other types of Cyp11a1-expressing cells. This explains why deletion of Cyp11a1 in T cells significantly reduced steroidogenesis and its effect. The presence of Cyp11a1^+^ non-T cells indicates that in a different context such as in different tumor types the steroidogenic population may vary. Depending upon the immunologic context and congressing of tumor associated cell types the intratumoral steroid pool may vary and play role beyond the immune suppression. For example, synthesis of estrogen, progesterone or androgen may have direct effect on cancer cell survival and proliferation, immunogenicity and that may impose another level of concern because breast, prostate and endometrial cancer are steroid sensitive and constitute a large fraction of all cancer occurrence ^47-49^. Previously it was reported that intratumoral *de novo* steroidogenesis promotes post-castration prostate cancer growth^50^. Though the role of immune cells in this context is unexplored. Future studies are essential to get detailed insight of immune cell-mediated intratumoral steroidogenesis in each tumor types.

The study strongly supports the concern raised by recent studies about proper use of steroids, particularly glucocorticoids, in solid tumors as a part of the disease management, specifically when we seek to boost anti-tumor immune response^51-58^. Glucocorticoids have been used in clinical oncology for over a half a century, and at present these are routinely used to alleviate edema in patients with intracranial lesions and are first-line agents to suppress immune-related adverse events that arise with the immunotherapies. Therefore, further studies are required to evaluate the use of synthetic steroids when the tumors itself induces glucocorticoid synthesis. Reduction of intratumoral steroidogenesis may open a window for immunotherapies to work better at lower doses.

Similar experimental approaches can in future provide in depth mechanistic insights into other extraglandular (local) steroidogenic cell types such as adipose cells, neuron, osteoblasts, astrocytes, microglia, skin, trophoblast and thymic epithelial cells. Our *Cyp11a1*-mCherry reporter mouse line can be used as a discovery tool to identify new steroidogenic cell types in tolerogenic physiological conditions such as pregnancy, mucosal tolerance, and inflammatory and immunopathological conditions. Their functional role can be dissected by using *Cyp11a1* cKO mice using tissue specific Cre-drivers. Nevertheless, cancer is a proven context where tumors hijack this pathway for immune evasion. Further studies in diverse physiological scenarios would be of great interest to more broadly understand the role of immune cell *de novo* steroidogenesis in regulating inflammation and immunity.

## Materials and Methods

### Mice

The care and use of all mice in this study were in accordance with the UK Animals in Science Regulation Unit’s Code of Practice for the Housing and Care of Animals Bred, Supplied or Used for Scientific Purposes, the Animals (Scientific Procedures) Act 1986 Amendment Regulations 2012. All procedures were performed under a UK Home Office Project license (PPL 80/2574 or PPL P8837835 or PPL P6B8058B0), which was reviewed and approved by the local institute’s Animal Welfare and Ethical Review Body.

### Generation of a Cyp11a1-mCherry reporter mouse line

*Targeting Description:* Using CRISPR_Cas9 (Clustered Regularly Interspaced Short Palindromic Repeats) ^59^ technology we introduced double strand DNA breaks 5’ and 3’ adjacent to the *Cyp11a1* termination codon in exon 9 to facilitate the introduction of our targeting construct. The 5’ and 3’ arms of homology were designed to remove the *Cyp11a1* termination codon and 100bp of the 3’ UTR immediately downstream and replace it with a minimal T2a self-cleavage peptide followed by the fluorescent marker mCherry.

### Cyp11a1 Guide RNA generation and ESC targeting

*sgRNA design and cloning:* Using the web-based tool designed by Hodgkins et al^60^, two sgRNAs were identified 5’ and 3’ adjacent to the Cyp11a1 termination codon. The guide sequences were ordered from Sigma Genosys as sense and antisense oligonucleotides, and annealed before individually cloning into the human U6 (hU6) expression plasmid (kind gift from Sebastian Gerety).

#### Targeted ES cell generation

3×10^6^ C57Bl/6 JM8 ESC (kind gift from Bill Skarnes) were nucleofected with 2ugs *Cyp11a1* circular targeting construct, 1.5ugs of each hU6_sgRNAs and 3ugs a plasmid expressing human codon-optimised CAS9 driven by the CMV promotor (Addgene # 41815). 48 hours post transfection the media was changed for G418 selection media. Confirmation of the correct targeting events were confirmed by qPCR for loss of heterozygosity (LOH) and the presence of the selectable marker, and long range (LR) PCR and Sanger sequencing.

#### Breeding and removal of the Selectable marker

Following blastocyst injection and chimaera breeding three F1 Cyp11a1+/^mCherry-Neo^ mice were breed to pCAGGSs-Flpo (kind gift from Andrea Kranz)^21^ mice to remove the neomycin selectable marker to generate Cyp11a1^+/mCherry^.

### Generation of a Cyp11a1 cKO mice

*Cyp11a1*^*fl/fl*^ mice were generated by crossing Cyp11a1^tm1a(KOMP)Wtsi^ mice with a previously reported Flp-deleter (FlpO) line^21^. *Cyp11a1*^*fl/fl*^ mice were crossed with *Cd4-cre* mice to generate the *Cyp11a1* cKO mice.

### Syngeneic melanoma model

B16-F10 melanoma model: The C57BL/6 derived B16-F10 melanoma cell line was purchased from American Type Culture Collection (ATCC) and cultured in Dulbecco’s Modified Eagle medium (DMEM, Life Technologies), supplemented with Penstrep and 10% FBS. For the primary tumor growth assay, 2.5 x10^5^ B16-F10 cells were injected subcutaneously into the shoulders of either wild type (WT) C57BL/6 mice, *Cd4-Cre, Cyp11a1*^*fl/fl*^ or *Cd4-Cre;Cyp11a1*^*fl/fl*^ mice. After 11 or 12 days animals were sacrificed and tissues collected for analysis. For the experimental metastasis assay, 5×10^5^ B16-F10 cells in a volume of 0.1 ml PBS were injected intra-venously into the tail vein. After ten days (±1 day) the mice were sacrificed *via* cervical dislocation, and their lungs removed and rinsed in phosphate buffered saline. The number of B16-F10 colonies on all five lobes of the lung were counted macroscopically.

Orthotopic EO771 (Synonym E0771) breast cancer model: The EO771 breast cancer cell line was purchased from CH3 BioSystems and cultured in RPMI (Sigma) supplemented with 10% FBS, 1% PS and 10mM HEPES. 2.5 x10^5^ cells were injected into the 4th inguinal mammary fat pad of Cyp11a1-mCherry reporter mice. Tumors and inguinal lymph node were collected after 15 days of tumor development.

Lung colonization of EO771 cells: EO771 cells were kindly gifted by Robin Anderson. X 10^5^ cells were injected intra-venously in to the tail vein. After ten days (±1 day) the mice were sacrificed via cervical dislocation, and their lungs were removed, ringed in PBS and fixed in. The fixed lungs were sectioned and stained with Hematoxylin and Eosin using professional service. The microscopic image were captured in Olympus with 10X magnification and area of the lung tumours were measured by ImageJ.

### Tumor Tissue Processing

Tumors were mechanically dissociated and digested in 1mg/ml collagenase D (Roche), 1mg/ml collagenase A (Roche) and 0.4mg/ml DNase I (Sigma) in IMDM media containing 10% FBS, at 37°C for 40 mins. EDTA was added to all samples to neutralize collagenase activity (final concentration 5mM) and digested tissues were passed through 70µm filters (Falcon).

### Cell sorting

Once processed, single cell suspension tumour samples were incubated with a fixable fluorescent viability stain (Life Technologies) for 20mins (diluted 1:1000 in PBS) prior to incubation with conjugated primary antibodies for 30 mins at 4°C. Antibodies were diluted in PBS 0.5% BSA. Stained samples were sorted, using the MoFlo XDP or BD Influx cytometer system.

### B16-F10 and T cell co-culture assay

Negatively purified splenic naïve CD4^+^ T cells from Cyp11a1-mCherry mice were cultured in presence of B16-F10 cell with or without TCR activation and analyzed by flow cytometry to detect Cyp11a1-mCherry expression. Naïve T cell purification and activation condition is described previously^61^. Briefly, naïve splenic cells were activated in anti-CD3e and anti-CD28 coated plate in presence of IL2 but in absence anti-IL4 neutralizing antibody.

### Quantitative PCR (qPCR)

Tumor infiltrating macrophages (Lin^−^CD11b^+^) and CD8^+^ T cells were purified by cell sorting. We used the Cells-to-C_T_ kit (Invitrogen/Thermofisher Scientific) and followed SYBR Green format according to manufacturer’s instructions. 2µl of cDNA was used in 12µl qPCR reactions with appropriate primers and SYBR Green PCR Master Mix (Applied Biosystems). Data were analyzed by ddCT method. Experiments were performed 3 times and data represent mean values ± standard deviation. The primer list is provided below:

*Arg1*: F- ATGGAAGAGACCTTCAGCTAC

R- GCTGTCTTCCCAAGAGTTGGG

*Tgfβ1*: F- TGACGTCACTGGAGTTGTACGG

R- GGTTCATGTCATGGATGGTGC

*Ifnγ*: F- ACAATGAACGCTACACACTGC

R- CTTCCACATCTATGCCACTTGAG;

*Tnfα*: F- CATCTTCTCAAAATTCGAGTGACAA

R- TGGGAGTAGACAAGGTACAACCC

*Gapdh*: F- ACCACAGTCCATGCCATCAC

R- GCCTGCTTCACCACCTTC

*Rplp0*: F: CACTGGTCTAGGACCCGAGAA

R: GGTGCCTCTGGAGATTTTCG

### Flow cytometry

We followed eBioscience surface staining, intracellular cytotoplasmic protein staining (for cytokines) protocols. Briefly, single cell suspension was stained with Live/Dead Fixable Dead cell stain kit (Molecular Probes/ Thermo Fisher) and blocked by purified rat anti-mouse CD16/CD32 purchased from BD Bioscience and eBioscience. Surface staining was performed in flow cytometry staining buffer (eBioscience) or in PBS containing 3% FCS at 4°C. For intracellular cytokine staining cells were fixed by eBioscience IC Fixation buffer and permeabililzed by eBioscience permeabilization buffer. Cells were stained in 1x permeabilization buffer with fluorescent dye-conjugated antibodies. After staining cells were washed with flow cytometry staining buffer (eBioscience) or 3% PBS-FCS, and were analyzed by flow cytometer Fortessa (BD Biosciences) using FACSDiva. The data were analyzed by FlowJo software. Antibodies used in flow cytometry were: CD4 (RM4-5 or GK1.5), CD8a (53-6.7), CD3e (145-2c11), CD45 (30F11), CD44 (IM7), CD25 (PC61), B220 (Ra3-6b2), Cyp11a1 (C-16, unconjugated, Santa Cruz; Fluorescent dye conjugated anti-goat secondary was used for staining), Ly6G (1A8), Ly6G/Ly6C (Gr-1) (RB6-8C5), Ly6C (HK1.4), CD11b (M1/70), CD11c (N418), CD19 (1D3), NK1.1(Pk136), Ter119 (TER119), PD-1 (J43), TIGIT (1G9), CD107a/LAMP1(1D4B). All antibodies were purchased from eBioscience, BD Bioscience or Biolegend.

### Western Blot Antibodies

Anti-CYP11A1 (Santa Cruz Biotechnology, C-16) and anti-TBP (Abcam) were used.

### Quantitative ELISA

CD45^+^ leukocytes were purified from B16-F10 tumor masses and lungs, of mice that had been tail vein administered B16-F0 cells, and seeded at equal density in IMDM medium supplemented with 10% charcoal stripped fetal bovine serum (Life Technologies, Invitrogen) for 24 hrs. Pregnenolone concentrations of the culture supernatants were quantified using pregnenolone ELISA kit (Abnova) and corticosteroids ELISA (Thermofisher) kit following manufacturers’ instruction. Absorbance was measured at 450 nm, and data were analyzed in GraphPad Prism 5.

### B16-F10 and T cell co-culture assay

Negatively purified splenic naïve CD4^+^ T cells from Cyp11a1-mCherry mice were cultured in presence of B16-F10 cell with or without TCR activation and analyzed by flow cytometry to detect Cyp11a1-mCherry expression.

## Acknowledgements

We would like to thank Ana C. Anderson and Rahul Roychoudhuri for their valuable comments on the manuscript and useful discussions; Jana Eliasova and Sarah Davidson for her help with diagram illustration; Bee Ling Ng, Chris Hall, Sam Thompson and Jennie Graham for help with flow cytometry and cell sorting; Research Support Facility, WSI, for their technical help and animal husbandry. EO771 cells were kindly gifted by Robin Anderson.

## Author contribution

BM: Led and managed the project, generated hypothesis, designed and performed experiments, analyzed data, and wrote manuscript JP: Performed experiments, analyzed data and helped in genetically modified mouse generation. LvdW: Performed B16-F10 and EO771 pulmonary metastasis experiments, analyzed data. AR: Performed B16-F10 subcutaneous tumors experiments, analyzed data. GK, NAF and KK: Analyzed publicly available gene expression datasets to confirm human tumor expression of steroidogenic genes. LSC: Helped in identification and quantification of tumor spots in EO771 lung metastasis. ER and GD: Helped in generation of *Cyp11a*-mCherry and *Cyp111a1*^*fl/fl*^ mouse model. IW: Built the *Cyp11a1*-mCherry targeting construct. KO: Helped in designing experiments, writing manuscript, critical comments and supervision. DJA: Conducted pulmonary metastasis experiments. JS: Conducted B16-F10 subcutaneous experiments under her PPL and supervised the study. SAT: Supervised the study. All authors commented on and approved of the draft manuscript before submission.

## REFERENCES

1 Miller, W. L. & Auchus, R. J. The molecular biology, biochemistry, and physiology of human steroidogenesis and its disorders. Endocr Rev 32, 81–151, doi:10.1210/er.2010-0013 (2011).

2 Miller, W. L. Steroidogenesis: Unanswered Questions. Trends in endocrinology and metabolism: TEM 28, 771–793, doi:10.1016/j.tem.2017.09.002 (2017).

3 Belelli, D. & Lambert, J. J. Neurosteroids: endogenous regulators of the GABA(A) receptor. Nature reviews. Neuroscience 6, 565–575, doi:10.1038/nrn1703 (2005).

4 Hosie, A. M., Wilkins, M. E., da Silva, H. M. & Smart, T. G. Endogenous neurosteroids regulate GABAA receptors through two discrete transmembrane sites. Nature 444, 486–489, doi:10.1038/nature05324 (2006).

5 Slominski, A. et al. Steroidogenesis in the skin: implications for local immune functions. The Journal of steroid biochemistry and molecular biology 137, 107–123, doi:10.1016/j.jsbmb.2013.02.006 (2013).

6 Vacchio, M. S., Papadopoulos, V. & Ashwell, J. D. Steroid production in the thymus: implications for thymocyte selection. The Journal of experimental medicine 179, 1835–1846 (1994).

7 Li, J., Papadopoulos, V. & Vihma, V. Steroid biosynthesis in adipose tissue. Steroids 103, 89–104, doi:10.1016/j.steroids.2015.03.016 (2015).

8 Cima, I. et al. Intestinal epithelial cells synthesize glucocorticoids and regulate T cell activation. The Journal of experimental medicine 200, 1635–1646, doi:10.1084/jem.20031958 (2004).

9 Hostettler, N. et al. Local glucocorticoid production in the mouse lung is induced by immune cell stimulation. Allergy 67, 227–234, doi:10.1111/j.1398-9995.2011.02749.x (2012).

10 Hanahan, D. & Weinberg, R. A. Hallmarks of cancer: the next generation. Cell 144, 646–674, doi:10.1016/j.cell.2011.02.013 (2011).

11 Vinay, D. S. et al. Immune evasion in cancer: Mechanistic basis and therapeutic strategies. Semin Cancer Biol 35 Suppl, S185–S198, doi:10.1016/j.semcancer.2015.03.004 (2015).

12 Beatty, G. L. & Gladney, W. L. Immune escape mechanisms as a guide for cancer immunotherapy. Clin Cancer Res 21, 687–692, doi:10.1158/1078-0432.CCR-14-1860 (2015).

13 DeNardo, D. G. et al. CD4(+) T cells regulate pulmonary metastasis of mammary carcinomas by enhancing protumor properties of macrophages. Cancer Cell 16, 91–102, doi:10.1016/j.ccr.2009.06.018 (2009).

14 Kobayashi, M., Kobayashi, H., Pollard, R. B. & Suzuki, F. A pathogenic role of Th2 cells and their cytokine products on the pulmonary metastasis of murine B16 melanoma. J Immunol 160, 5869–5873 (1998).

15 De Monte, L. et al. Intratumor T helper type 2 cell infiltrate correlates with cancer-associated fibroblast thymic stromal lymphopoietin production and reduced survival in pancreatic cancer. J Exp Med 208, 469–478, doi:10.1084/jem.20101876 (2011).

16 Hanahan, D. & Coussens, L. M. Accessories to the crime: functions of cells recruited to the tumor microenvironment. Cancer Cell 21, 309–322, doi:10.1016/j.ccr.2012.02.022 (2012).

17 Cain, D. W. & Cidlowski, J. A. Immune regulation by glucocorticoids. Nature reviews. Immunology 17, 233–247, doi:10.1038/nri.2017.1 (2017).

18 Klein, S. L. & Flanagan, K. L. Sex differences in immune responses. Nature reviews. Immunology 16, 626–638, doi:10.1038/nri.2016.90 (2016).

19 Mahata, B. et al. Single-cell RNA sequencing reveals T helper cells synthesizing steroids de novo to contribute to immune homeostasis. Cell Rep 7, 1130–1142, doi:10.1016/j.celrep.2014.04.011 (2014).

20 Skarnes, W. C. et al. A conditional knockout resource for the genome-wide study of mouse gene function. Nature 474, 337–342, doi:10.1038/nature10163 (2011).

21 Kranz, A. et al. An improved Flp deleter mouse in C57Bl/6 based on Flpo recombinase. Genesis 48, 512–520, doi:10.1002/dvg.20641 (2010).

22 Mellman, I., Coukos, G. & Dranoff, G. Cancer immunotherapy comes of age. Nature 480, 480–489, doi:10.1038/nature10673 (2011).

23 Landskron, G., De la Fuente, M., Thuwajit, P., Thuwajit, C. & Hermoso, M. A. Chronic inflammation and cytokines in the tumor microenvironment. Journal of immunology research 2014, 149185, doi:10.1155/2014/149185 (2014).

24 Papatheodorou, I. et al. Expression Atlas: gene and protein expression across multiple studies and organisms. Nucleic acids research 46, D246–D251, doi:10.1093/nar/gkx1158 (2018).

25 Curran, M. A., Montalvo, W., Yagita, H. & Allison, J. P. PD-1 and CTLA-4 combination blockade expands infiltrating T cells and reduces regulatory T and myeloid cells within B16 melanoma tumors. Proceedings of the National Academy of Sciences of the United States of America 107, 4275–4280, doi:10.1073/pnas.0915174107 (2010).

26 De Henau, O. et al. Overcoming resistance to checkpoint blockade therapy by targeting PI3Kgamma in myeloid cells. Nature 539, 443–447, doi:10.1038/nature20554 (2016).

27 Zamarin, D. et al. Intratumoral modulation of the inducible co-stimulator ICOS by recombinant oncolytic virus promotes systemic anti-tumour immunity. Nature communications 8, 14340, doi:10.1038/ncomms14340 (2017).

28 Kisielow, J., Obermair, F. J. & Kopf, M. Deciphering CD4(+) T cell specificity using novel MHC-TCR chimeric receptors. Nat Immunol 20, 652–662, doi:10.1038/s41590-019-0335-z (2019).

29 Casey, A. E., Laster, W. R., Jr. & Ross, G. L. Sustained enhanced growth of carcinoma EO771 in C57 black mice. Proc Soc Exp Biol Med 77, 358–362, doi:10.3181/00379727-77-18779 (1951).

30 Johnstone, C. N. et al. Functional and molecular characterisation of EO771.LMB tumours, a new C57BL/6-mouse-derived model of spontaneously metastatic mammary cancer. Dis Model Mech 8, 237–251, doi:10.1242/dmm.017830 (2015).

31 van der Weyden, L. et al. Genome-wide in vivo screen identifies novel host regulators of metastatic colonization. Nature 541, 233–236, doi:10.1038/nature20792 (2017).

32 Martinez, F. O. & Gordon, S. The M1 and M2 paradigm of macrophage activation: time for reassessment. F1000Prime Rep 6, 13, doi:10.12703/P6-13 (2014).

33 Roszer, T. Understanding the Mysterious M2 Macrophage through Activation Markers and Effector Mechanisms. Mediators Inflamm 2015, 816460, doi:10.1155/2015/816460 (2015).

34 Desbarats, J., You-Ten, K. E. & Lapp, W. S. Levels of p56lck and p59fyn are reduced by a glucocorticoid-dependent mechanism in graft-versus-host reaction-induced T cell anergy. Cell Immunol 163, 10–18, doi:10.1006/cimm.1995.1093 (1995).

35 Van Laethem, F. et al. Glucocorticoids attenuate T cell receptor signaling. J Exp Med 193, 803–814, doi:10.1084/jem.193.7.803 (2001).

36 Heijink, I. H. & Van Oosterhout, A. J. Strategies for targeting T-cells in allergic diseases and asthma. Pharmacol Ther 112, 489–500, doi:10.1016/j.pharmthera.2006.05.005 (2006).

37 Gordon, J. R., Ma, Y., Churchman, L., Gordon, S. A. & Dawicki, W. Regulatory dendritic cells for immunotherapy in immunologic diseases. Front Immunol 5, 7, doi:10.3389/fimmu.2014.00007 (2014).

38 Piemonti, L. et al. Glucocorticoids affect human dendritic cell differentiation and maturation. J Immunol 162, 6473–6481 (1999).

39 Woltman, A. M. et al. The effect of calcineurin inhibitors and corticosteroids on the differentiation of human dendritic cells. Eur J Immunol 30, 1807–1812, doi:10.1002/1521-4141(200007)30:7<1807::AID-IMMU1807>3.0.CO;2-N (2000).

40 Chen, X., Oppenheim, J. J., Winkler-Pickett, R. T., Ortaldo, J. R. & Howard, O. M. Glucocorticoid amplifies IL-2-dependent expansion of functional FoxP3(+)CD4(+)CD25(+) T regulatory cells in vivo and enhances their capacity to suppress EAE. Eur J Immunol 36, 2139–2149, doi:10.1002/eji.200635873 (2006).

41 Hu, Y. et al. Function of regulatory T-cells improved by dexamethasone in Graves’ disease. Eur J Endocrinol 166, 641–646, doi:10.1530/EJE-11-0879 (2012).

42 Suarez, A., Lopez, P., Gomez, J. & Gutierrez, C. Enrichment of CD4+ CD25high T cell population in patients with systemic lupus erythematosus treated with glucocorticoids. Ann Rheum Dis 65, 1512–1517, doi:10.1136/ard.2005.049924 (2006).

43 Sakuishi, K. et al. Targeting Tim-3 and PD-1 pathways to reverse T cell exhaustion and restore anti-tumor immunity. The Journal of experimental medicine 207, 2187–2194, doi:10.1084/jem.20100643 (2010).

44 Anderson, A. C., Joller, N. & Kuchroo, V. K. Lag-3, Tim-3, and TIGIT: Co-inhibitory Receptors with Specialized Functions in Immune Regulation. Immunity 44, 989–1004, doi:10.1016/j.immuni.2016.05.001 (2016).

45 Chihara, N. et al. Induction and transcriptional regulation of the co-inhibitory gene module in T cells. Nature 558, 454–459, doi:10.1038/s41586-018-0206-z (2018).

46 Rhen, T. & Cidlowski, J. A. Antiinflammatory action of glucocorticoids--new mechanisms for old drugs. N Engl J Med 353, 1711–1723, doi:10.1056/NEJMra050541 (2005).

47 DeVita, V. T., Lawrence, T. S. & Rosenberg, S. A. DeVita, Hellman, and Rosenberg’s cancer : principles & practice of oncology. 11th edition. edn, (Wolters Kluwer, 2019).

48 Risbridger, G. P., Davis, I. D., Birrell, S. N. & Tilley, W. D. Breast and prostate cancer: more similar than different. Nat Rev Cancer 10, 205–212, doi:10.1038/nrc2795 (2010).

49 Salmon, H., Remark, R., Gnjatic, S. & Merad, M. Host tissue determinants of tumour immunity. Nat Rev Cancer 19, 215–227, doi:10.1038/s41568-019-0125-9 (2019).

50 Locke, J. A. et al. Androgen levels increase by intratumoral de novo steroidogenesis during progression of castration-resistant prostate cancer. Cancer Res 68, 6407–6415, doi:10.1158/0008-5472.CAN-07-5997 (2008).

51 Flint, T. R. et al. Tumor-Induced IL-6 Reprograms Host Metabolism to Suppress Anti-tumor Immunity. Cell Metab 24, 672–684, doi:10.1016/j.cmet.2016.10.010 (2016).

52 Connell, C. M. et al. Cancer immunotherapy trial registrations increase exponentially but chronic immunosuppressive glucocorticoid therapy may compromise outcomes. Ann Oncol 28, 1678–1679, doi:10.1093/annonc/mdx181 (2017).

53 Della Corte, C. M. & Morgillo, F. Early use of steroids affects immune cells and impairs immunotherapy efficacy. ESMO Open 4, e000477, doi:10.1136/esmoopen-2018-000477 (2019).

54 Draghi, A. et al. Differential effects of corticosteroids and anti-TNF on tumor-specific immune responses: implications for the management of irAEs. Int J Cancer 145, 1408–1413, doi:10.1002/ijc.32080 (2019).

55 Obradovic, M. M. S. et al. Glucocorticoids promote breast cancer metastasis. Nature 567, 540–544, doi:10.1038/s41586-019-1019-4 (2019).

56 Giles, A. J. et al. Dexamethasone-induced immunosuppression: mechanisms and implications for immunotherapy. J Immunother Cancer 6, 51, doi:10.1186/s40425-018-0371-5 (2018).

57 Arbour, K. C. et al. Impact of Baseline Steroids on Efficacy of Programmed Cell Death-1 and Programmed Death-Ligand 1 Blockade in Patients With Non-Small-Cell Lung Cancer. J Clin Oncol 36, 2872–2878, doi:10.1200/JCO.2018.79.0006 (2018).

58 Tokunaga, A. et al. Selective inhibition of low-affinity memory CD8(+) T cells by corticosteroids. J Exp Med, doi:10.1084/jem.20190738 (2019).

59 Wiedenheft, B. et al. Structures of the RNA-guided surveillance complex from a bacterial immune system. Nature 477, 486–489, doi:10.1038/nature10402 (2011).

60 Hodgkins, A. et al. WGE: a CRISPR database for genome engineering. Bioinformatics 31, 3078–3080, doi:10.1093/bioinformatics/btv308 (2015).

61 Pramanik, J. et al. Genome-wide analyses reveal the IRE1a-XBP1 pathway promotes T helper cell differentiation by resolving secretory stress and accelerating proliferation. Genome Med 10, 76, doi:10.1186/s13073-018-0589-3 (2018).

