## Additional Data Figures for "Tumors induce *de novo* steroid biosynthesis in T cells to evade immunity"

**Extended Data Figure 1**

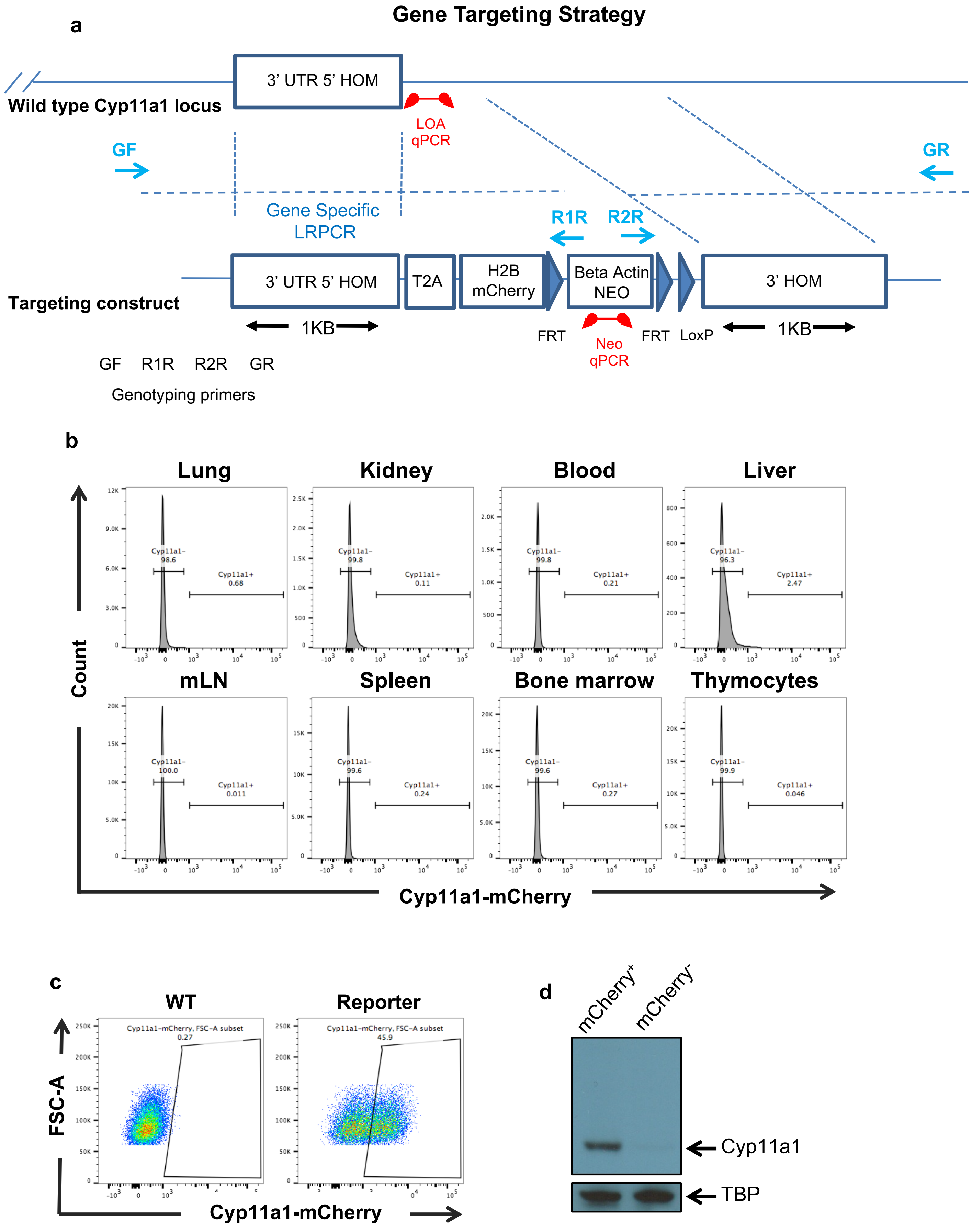

**Extended Data Figure 1. Generation of a Cyp11a1-mCherry reporter mouse line that reports Cyp11a1 expression accurately.**

a. Diagrammatic presentation of the targeting allele and strategy of Cyp11a1-mCherry mouse line generation.

b. Tissue expression of Cyp11a1-mCherry in naïve mice. Lung, kidney, blood, liver, mesenteric lymph node (mLN), spleen, bone marrow and thymus were harvested from Cyp11a1-mCherry reporter mice. All tissues were mechanically dissociated and for kidney, liver and lung were additionally enzymatically digested into single cell suspension and analysed by flow cytometry. Gating: All cells > Singlets > Live cells > Cyp11a1-mCherry. Representative data of three independent repeats (n=9).

c. Splenic naïve $CD4^+$ T cells from wild type (WT) and Cyp11a1-mCherry reporter mice were purified by negative selection using MACS, activated in vitro under Th2 differentiation condition, and analysed by flow cytometry. Gating: All cells > Singlets > Live cells > Cyp11a1-mCherry. Representative of more than six independent experiments.

d. Splenic naïve $CD4^+$ T cells from Cyp11a1-mCherry reporter mice were purified by negative selection, activated in vitro under Th1 and Th2 differentiation condition. Differentiated Th1 and Th2 cells were mixed together and mCherry+ and mCherry- cells were sorted by cell sorter. Cyp11a1 expression was analyzed by western blotting. Representative of three independent experiments (n=3).

### Extended Data Figure 2

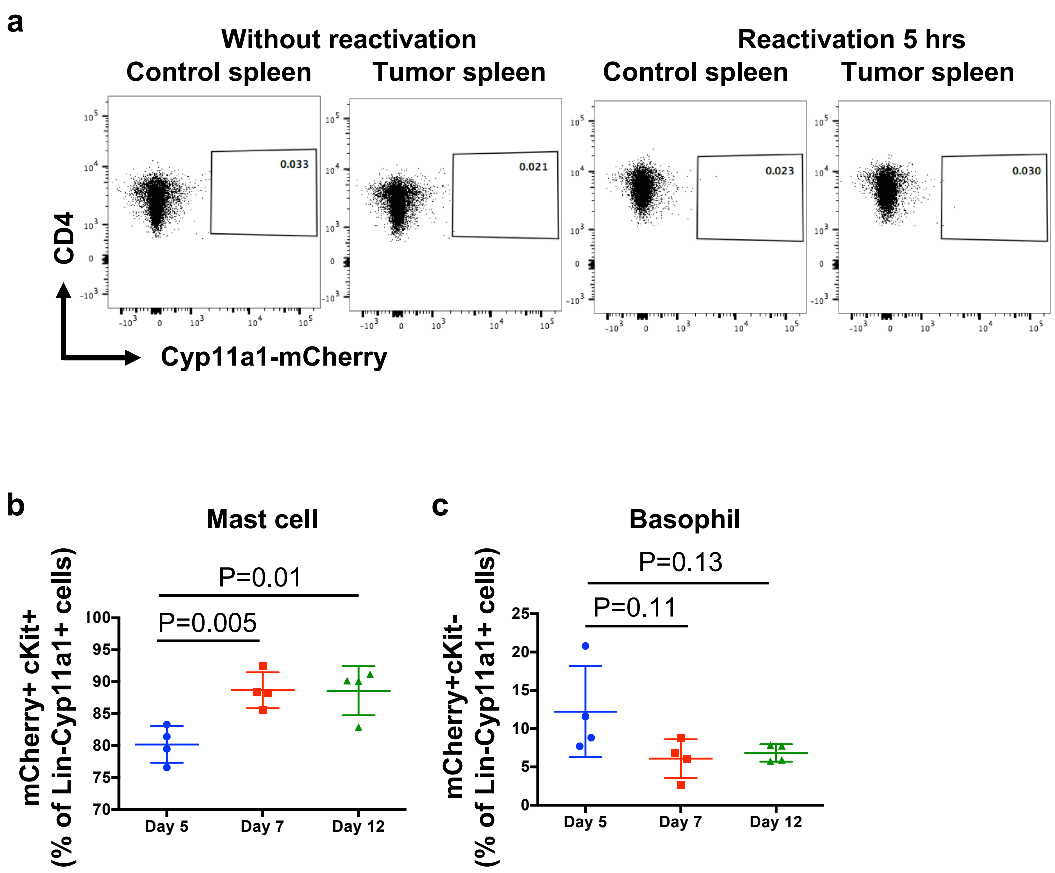

#### Extended Data Figure 2

d

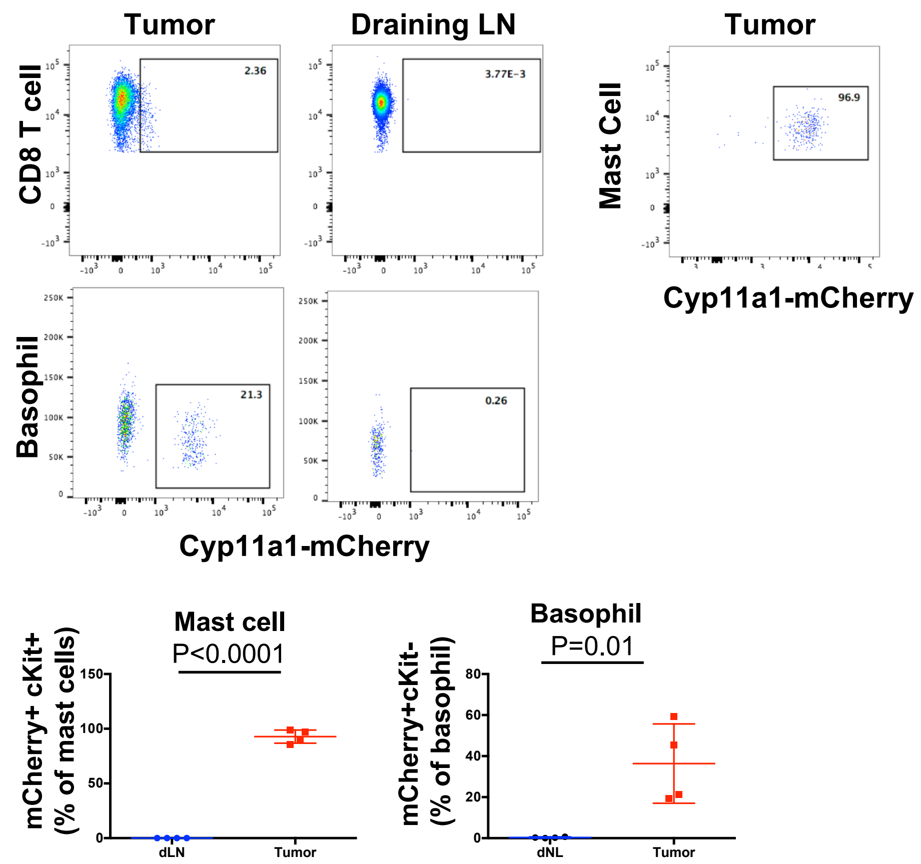


e

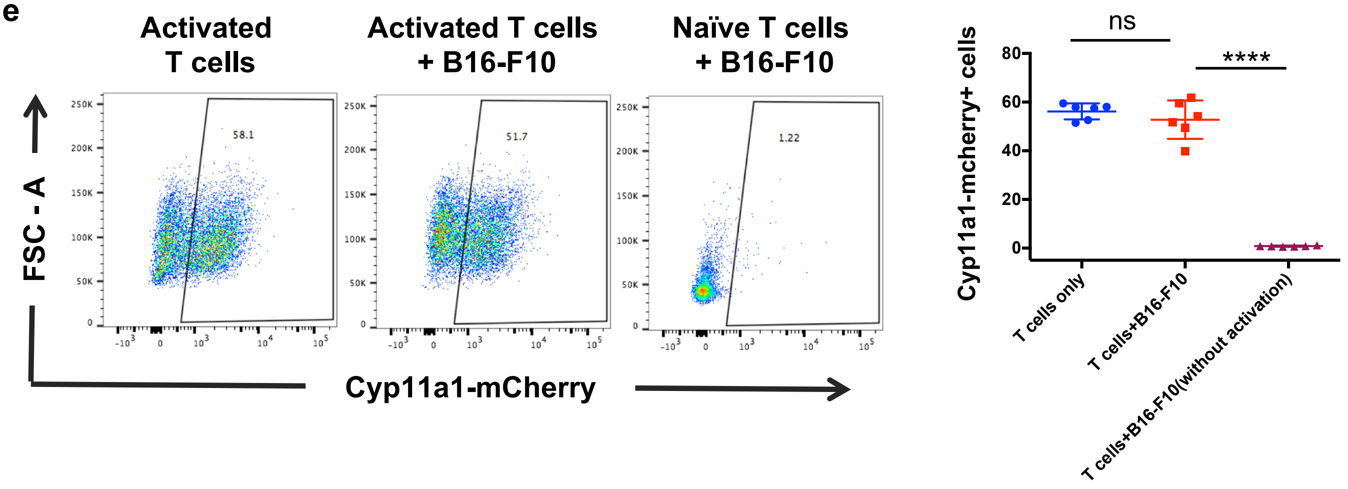

#### Extended Data Figure 2

**f**

##### Expression in tumor tissue

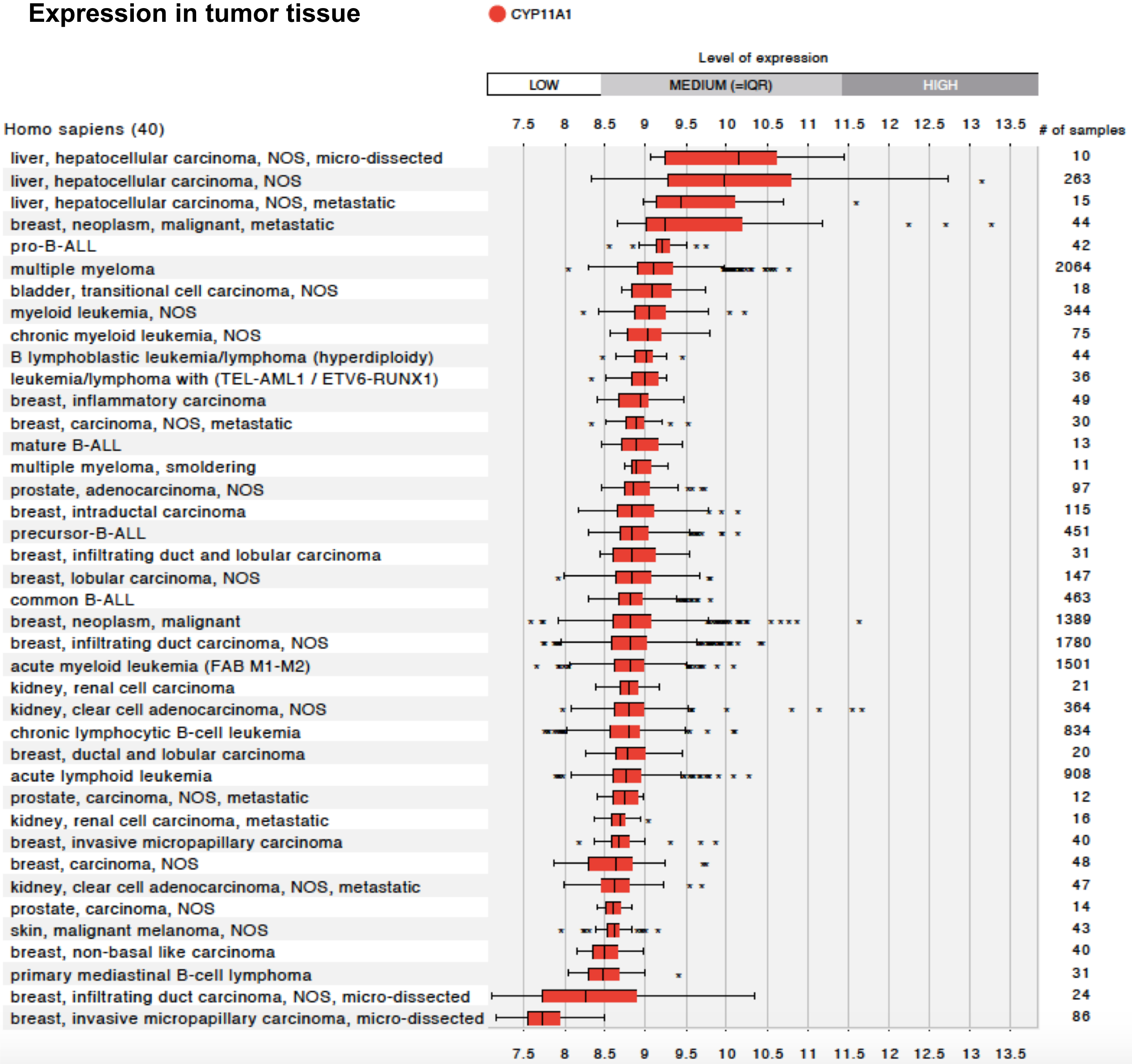


##### Expression in cancer cell lines

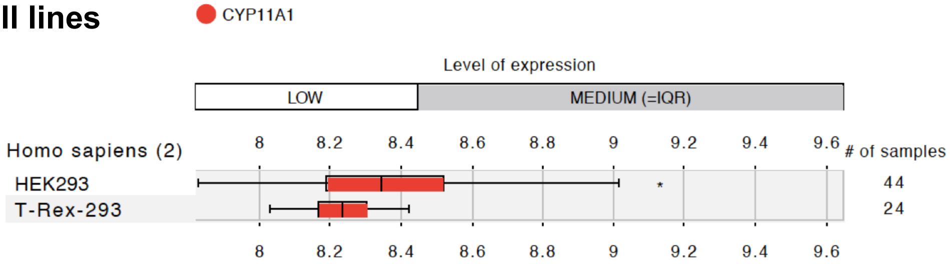

### Extended Data Figure 2

g

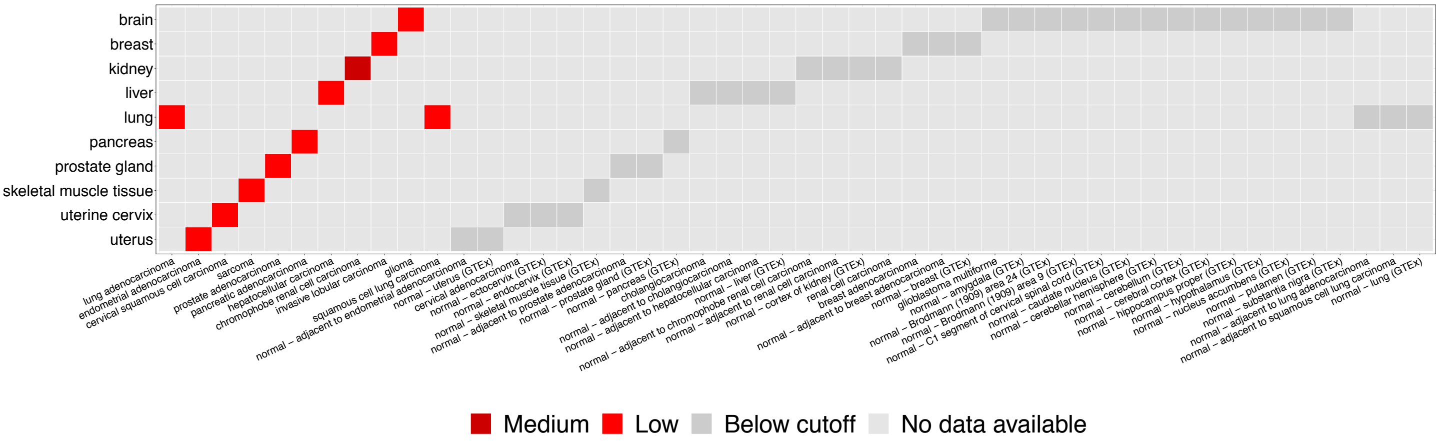


h

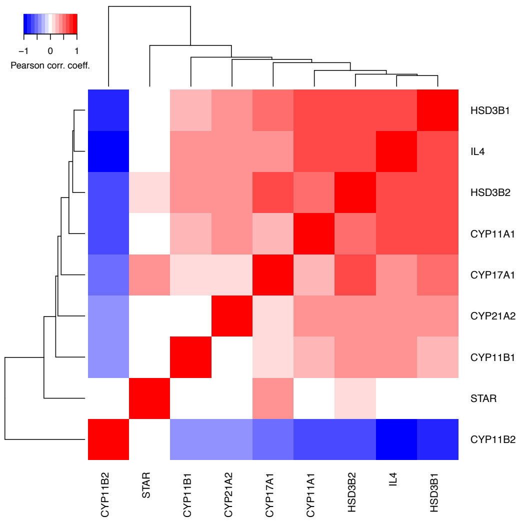

**Extended Data Figure 2. In vivo induction of Cyp11a1 in cancer**

**a-d.** Induction of Cyp11a1 in mouse models of tumors.

**a.** Splenic T cells do not express Cyp11a1 even after restimulation by PMA and ionomycin. Representative FACS profile of Figure 2b.

**b-c.** *In vivo* intratumoral Cyp11a1 expression dynamics in mast cell and basophil. B16-F10 cells were injected subcutaneously into the shoulder region of Cyp11a1-mCherry reporter mice. After 5, 7 and 12 days tumor tissues were dissociated into single cell suspensions, and analyzed by flow cytometry to detect the mast cell ($CD45^{+}Lin^{-}Fcer1^{+}cKit^{+}$) (**b**), and basophils ($CD45^{+}Lin^{-}Fcer1^{+}cKit^{-}SiglecF^{-}$) (**c**).

**d.** Immune cell-mediated Cyp11a1 expression is conserved in EO771 orthotopic model of triple negative breast cancer. EO771 cells were injected into the mammary fat pad of Cyp11a1-mCherry reporter mice. After 15 days, tumor tissues and tumor draining LN were dissociated into single cell suspensions, and analyzed by flow cytometry to detect the Cyp11a1-mCherry expression in $CD4^{+}$ T cells ($CD45^{+}Lin^{-}CD4^{+}CD3e^{+}CD8^{-}$), $CD8^{+}$ T cells ($CD45^{+}Lin\text{-}CD8a^{+}CD3e^{+}CD4^{-}$), mast cell ($CD45^{+}Lin^{-}Fcer1^{+}cKit^{+}$), and basophils ($CD45^{+}Lin^{-}Fcer1^{+}cKit^{-}SiglecF^{-}$). Graphical representation of mast cell and basophils are shown in the bottom panel.

e. Cancer cells (B16-F10 melanoma) do not induce Cyp11a1 expression directly. Splenic naïve $CD4^{+}$ T cells from Cyp11a1-mCherry mice were cultured in the presence or absence of B16-F10 cells with or without TCR activation and analyzed by flow cytometry to detect Cyp11a1-mCherry expression. Representative FACS profiles are shown in the left panels and graphical presentation is shown in the right panel.

f-g. **Induction of steroidogenesis in human tumors.** Publicly available data sets (GEO, ArrayExpress and TCGA) were analyzed to check steroidogenic genes and cytokine genes expression and their correlation. Tumors of the steroidogenic tissues, such as adrenal glands and gonads, were excluded from the analysis.

f. CYP11A1 expression and upregulation in human tumor types. Expression range of CYP11A1 in human tumors are shown using Genevestigator tool, which collects data from GEO, ArrayExpress and TCGA. Studies with at least 10 samples are selected for visualizing the expression box plot (top panel). Expression values are scaled between the experiments to make the expression values comparable using standard normalization methods implemented by the tool Genevestigator. CYP11A1 expression in kidney cancer cell line has been shown in the bottom panel.

g. CYP11A1 expression in human tumor samples and corresponding normal tissues as revealed in the Pan-Cancer Analysis of Whole Genomes study. CYP11A1 expression was searched at Expression Atlas, EMBL-EBI. Data were downloaded and the figure was reconstructed according to expression rank excluding the tumors of steroidogenic tissues (i.e. Ovary).

j. Heatmap showing correlation of steroidogenic gene expression and IL4 expression (Raw data source: GEO: GSE19234).

### Extended Data Figure 3

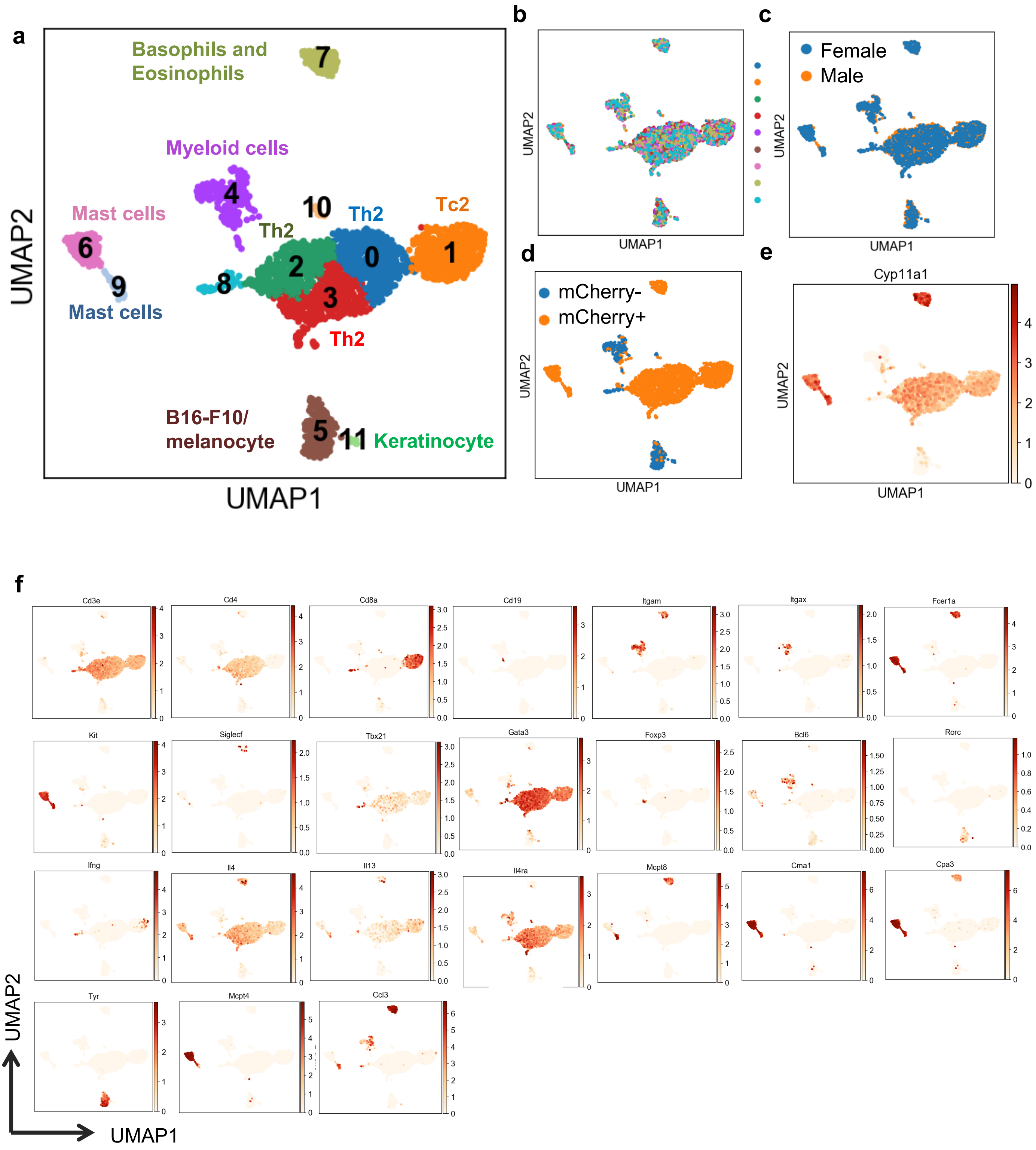

### Extended Data Figure 3

g

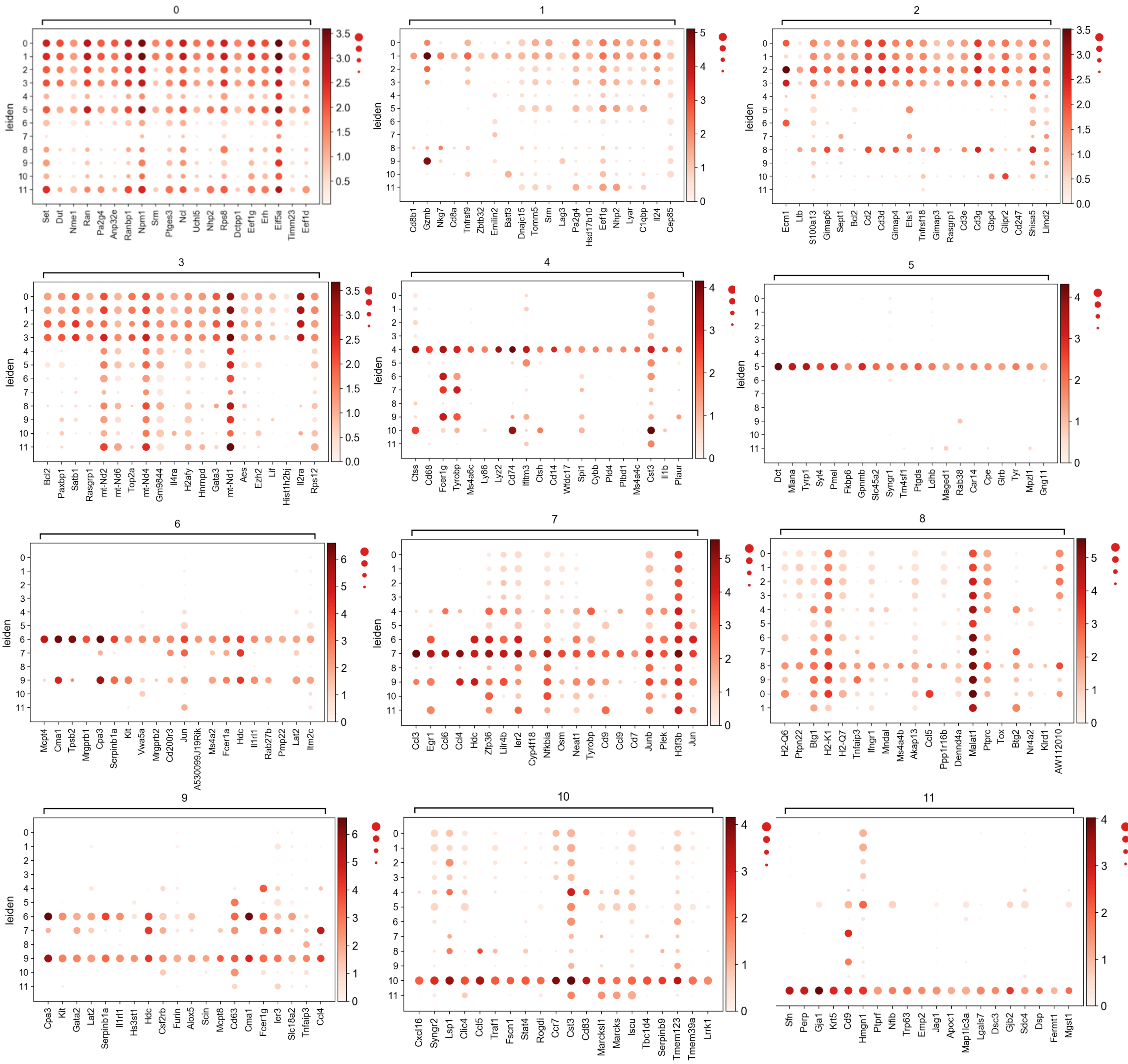


h

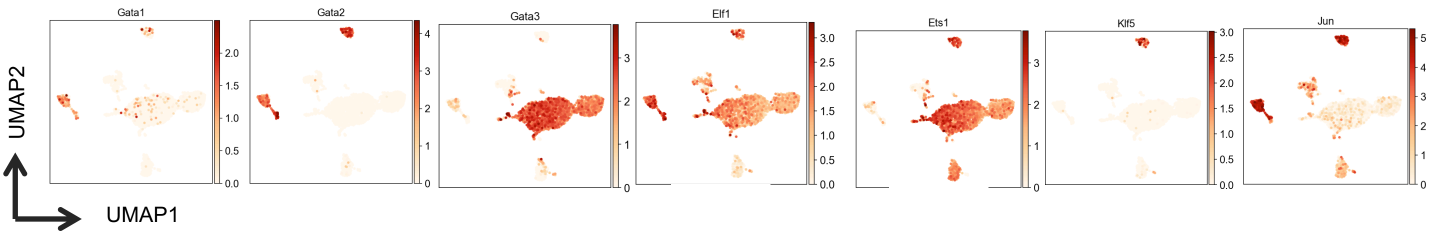

**Extended Data Figure 3**

**i**

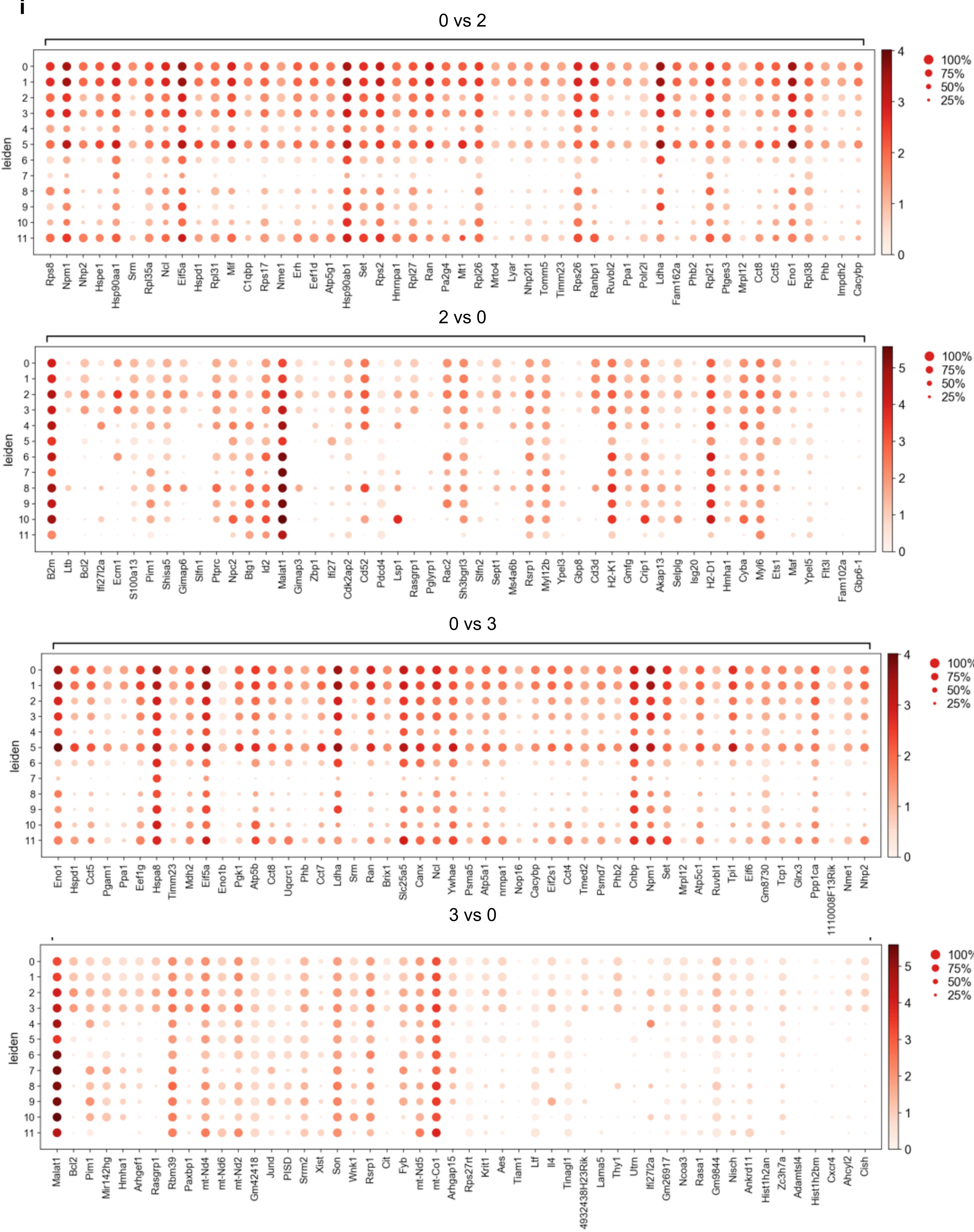

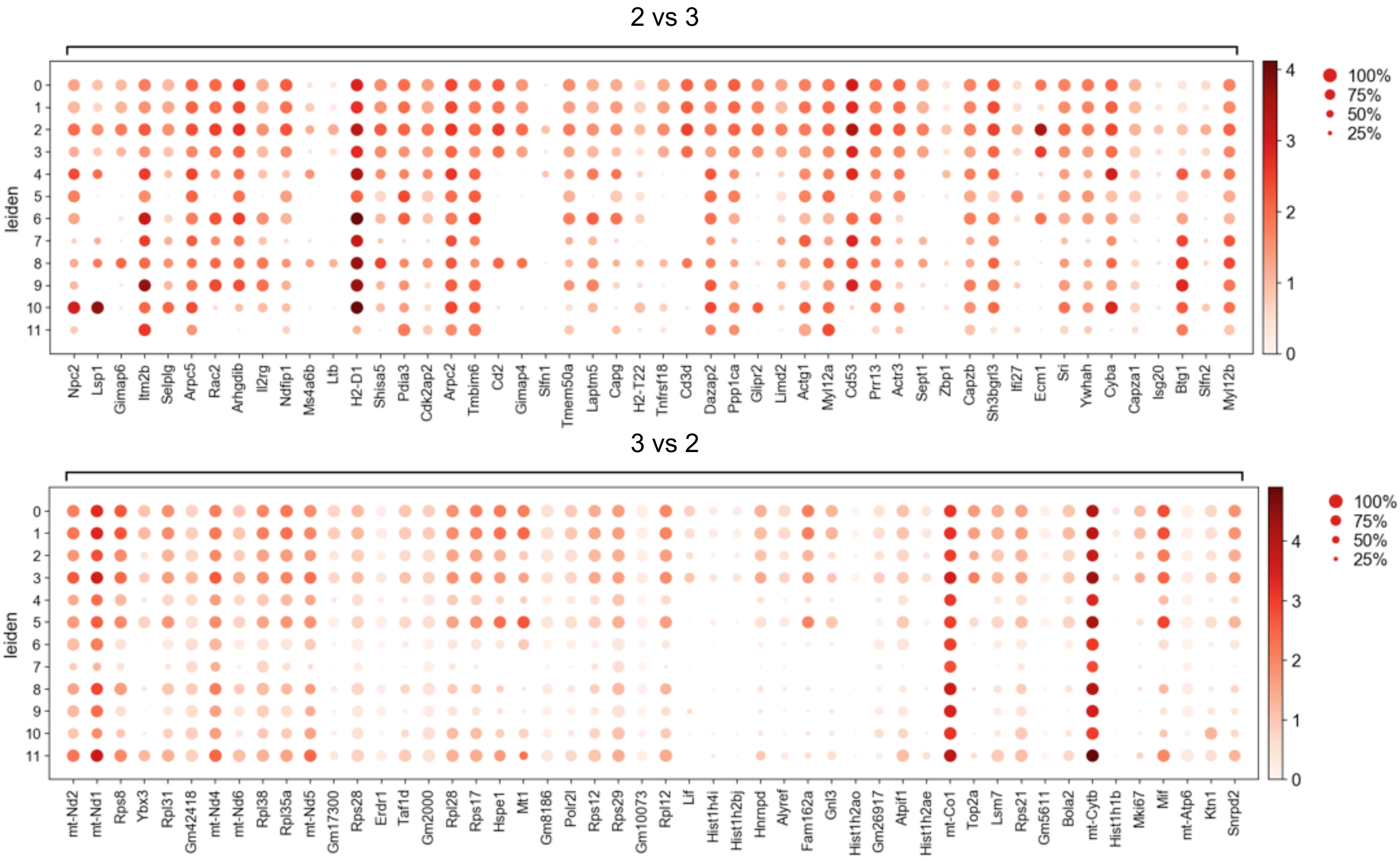
2 vs 3
leiden
0 1 2 3 4 5 6 7 8 9 10 11
Npc2 Lsp1 Gimap6 Itm2b Selplg Arpc5 Rac2 Arhgdib Il2rg Ndfip1 Ms4a6b Ltb H2-D1 Shisa5 Pdia3 Cdk2ap2 Arpc2 Tmbim6 Cd2 Gimap4 Slfn1 Tmem50a Laptm5 Capg H2-T22 Tnfrsf18 Cd3d Dazap2 Ppp1ca Glipr2 Limd2 Actg1 Myl12a Cd53 Prr13 Actr3 Sept1 Zbp1 Capzb Sh3bgrl3 Ifi27 Ecm1 Sri Ywhah Cyba Capza1 Isg20 Btg1 Slfn2 Myl12b
100%
75%
50%
25%
3 vs 2
mt-Nd2 mt-Nd1 Rps8 Ybx3 Rpl31 Gm42418 mt-Nd4 mt-Nd6 Rpl38 Rpl35a mt-Nd5 Gm17300 Rps28 Erdr1 Taf1d Gm2000 Rpl28 Rps17 Hspe1 Mt1 Gm8186 Polr2l Rps12 Rps29 Gm10073 Rpl12 Lif Hist1h4i Hist1h2bj Hnrnpd Alyref Fam162a Gnl3 Hist1h2ao Gm26917 Atpif1 Hist1h2ae mt-Co1 Top2a Lsm7 Rps21 Gm5611 Bola2 mt-Cytb Hist1h1b Mki67 Mif mt-Atp6 Ktn1 Snrpd2

**j**

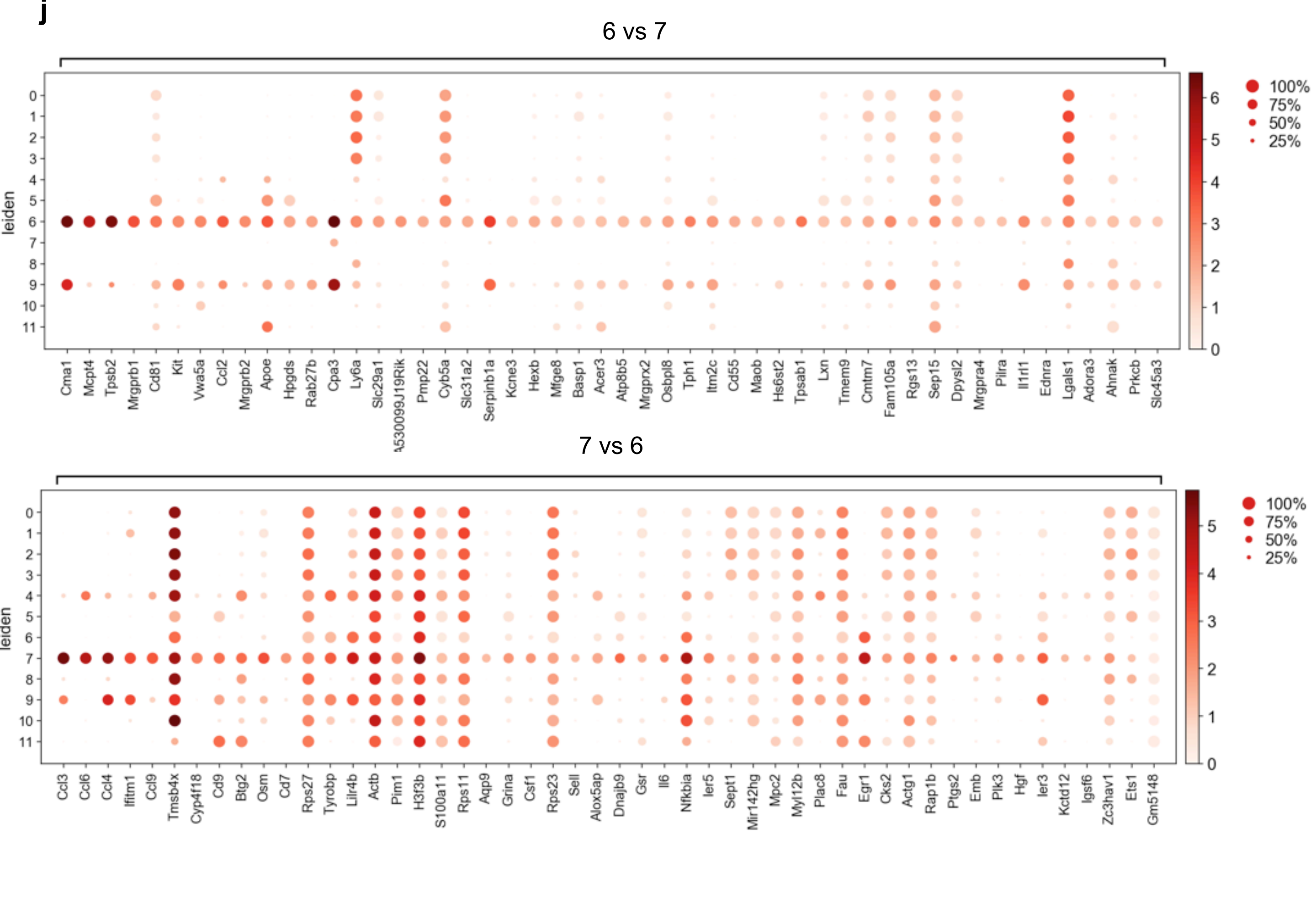
6 vs 7
leiden
0 1 2 3 4 5 6 7 8 9 10 11
Cma1 Mcpt4 Tpsb2 Mrgprb1 Cd81 Kit Vwa5a Ccl2 Mrgprb2 Apoe Hpgds Rab27b Cpa3 Ly6a Slc29a1 Pmp22 Cyb5a Slc31a2 Serpinb1a Kcne3 Hexb Mfge8 Basp1 Acer3 Atp8b5 Mrgprx2 Osbpl8 Tph1 Itm2c Cd55 Maob Hs6st2 Tpsab1 Lxn Tmem9 Cmtm7 Fam105a Rgs13 Sep15 Dpysl2 Mrgpra4 Pilra Il1rl1 Ednra Lgals1 Adora3 Ahnak Prkcb Slc45a3
100%
75%
50%
25%
7 vs 6
Ccl3 Ccl6 Ccl4 Ifitm1 Ccl9 Tmsb4x Cyp4f18 Cd9 Btg2 Osm Cd7 Rps27 Tyrobp Lilr4b Actb Pim1 H3f3b S100a11 Rps11 Aqp9 Grina Csf1 Rps23 Sell Alox5ap Dnajb9 Gsr Il6 Nfkbia Ier5 Sept1 Mir142hg Mpc2 Myl12b Plac8 Fau Egr1 Cks2 Actg1 Rap1b Ptgs2 Emb Plk3 Hgf Ier3 Kctd12 Igsf6 Zc3hav1 Ets1 Gm5148

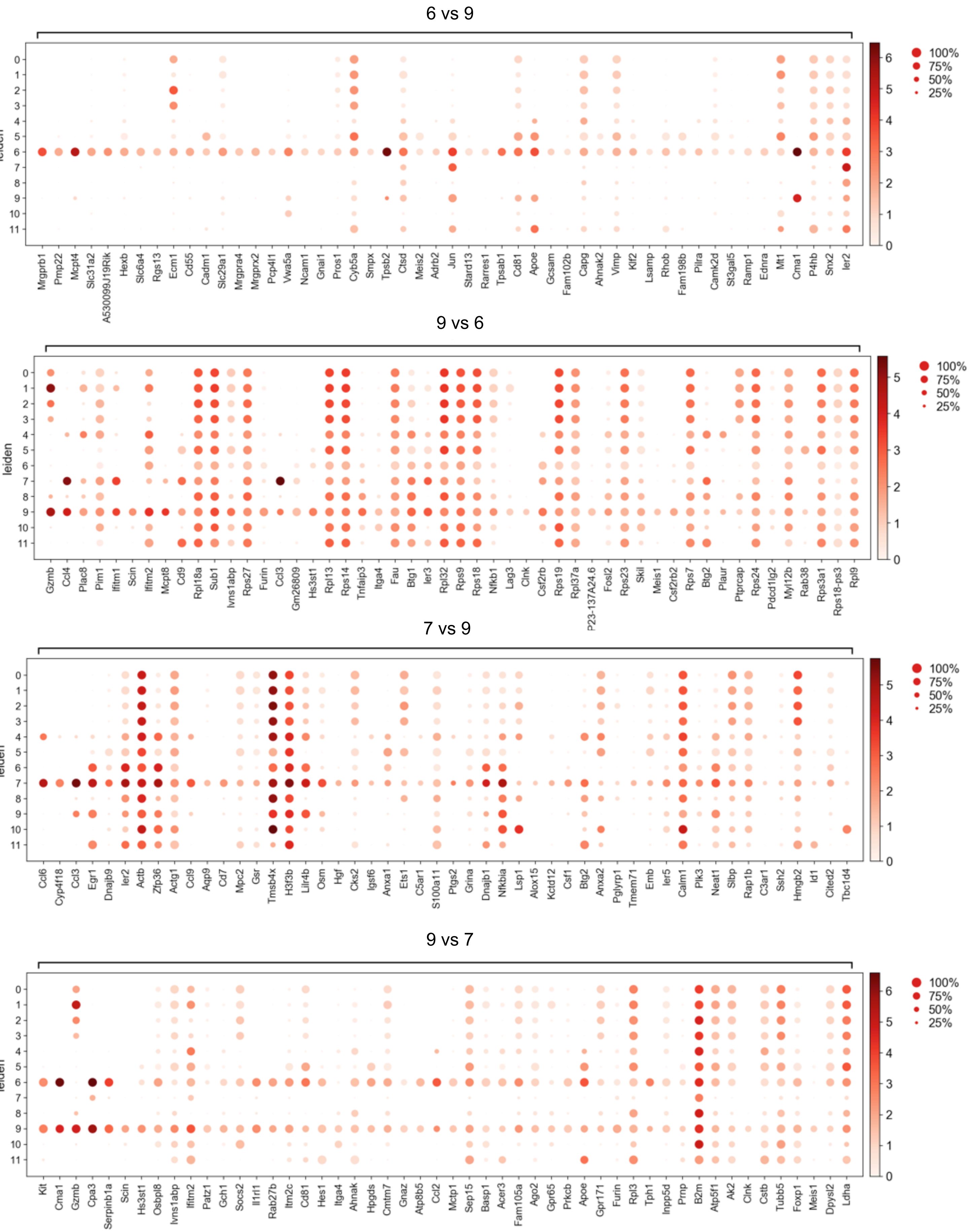
6 vs 9
leiden
0 1 2 3 4 5 6 7 8 9 10 11
Mrgprb1 Pmp22 Mcpt4 Slc31a2 A530099J19Rik Hexb Slc6a4 Rgs13 Ecm1 Cd55 Cadm1 Slc29a1 Mrgpra4 Mrgprx2 Pcp4l1 Vwa5a Ncam1 Gnai1 Pros1 Cyb5a Smpx Tpsb2 Ctsd Meis2 Adrb2 Jun Stard13 Rarres1 Tpsab1 Cd81 Apoe Gcsam Fam102b Capg Ahnak2 Vim Klf2 Lsamp Rhob Fam198b Pilra Camk2d St3gal5 Ramp1 Ednra Mt1 Cma1 P4hb Snx2 Ier2
0 1 2 3 4 5 6
100% 75% 50% 25%
9 vs 6
leiden
0 1 2 3 4 5 6 7 8 9 10 11
Gzmb Ccl4 Plac8 Pim1 Ifitm1 Scin Ifitm2 Mcpt8 Cd9 Rpl18a Sub1 Ivns1abp Rps27 Furin Ccl3 Gm26809 Hs3st1 Rpl13 Rps14 Tnfaip3 Itga4 Fau Btg1 Ier3 Rpl32 Rps9 Rps18 Nfkb1 Lag3 Clnk Csf2rb Rps19 Rpl37a P23-137A24.6 Fosl2 Rps23 Skil Meis1 Csf2rb2 Rps7 Btg2 Plaur Ptprcap Rps24 Pdcd1lg2 Myl12b Rab38 Rps3a1 Rps18-ps3 Rpl9
0 1 2 3 4 5
100% 75% 50% 25%
7 vs 9
leiden
0 1 2 3 4 5 6 7 8 9 10 11
Ccl6 Cyp4f18 Ccl3 Egr1 Dnajb9 Ier2 Actb Zfp36 Actg1 Ccl9 Aqp9 Cd7 Mpc2 Gsr Tmsb4x H3f3b Lilr4b Osm Hgf Cks2 Igsf6 Anxa1 Ets1 C5ar1 S100a11 Ptgs2 Grina Dnajb1 Nfkbia Lsp1 Alox15 Kctd12 Csf1 Btg2 Anxa2 Pglyrp1 Tmem71 Emb Ier5 Calm1 Plk3 Neat1 Slbp Rap1b C3ar1 Ssh2 Hmgb2 Id1 Cited2 Tbc1d4
0 1 2 3 4 5
100% 75% 50% 25%
9 vs 7
leiden
0 1 2 3 4 5 6 7 8 9 10 11
Kit Cma1 Gzmb Cpa3 Serpinb1a Scin Hs3st1 Osbpl8 Ivns1abp Ifitm2 Patz1 Gch1 Socs2 Il1rl1 Rab27b Itm2c Cd81 Hes1 Itga4 Ahnak Hpgds Cmtm7 Gnaz Atp8b5 Ccl2 Mctp1 Sep15 Basp1 Acer3 Fam105a Ago2 Gpr65 Prkcb Apoe Gpr171 Furin Rpl3 Tph1 Inpp5d Prnp B2m Atp5f1 Ak2 Clnk Cstb Tubb5 Foxp1 Meis1 Dpysl2 Ldha
0 1 2 3 4 5 6
100% 75% 50% 25%

### Extended Data Figure 3

**k**

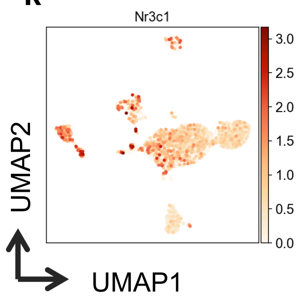

**Extended Data Figure 3. Single-cell transcriptomics revealed gene expression identity and gene expression pattern of intratumoral steroidogenic immune cells**

**a.** UMAP visualization of the tumor infiltrating cells with annotations of the clusters.

**b.** The clusters were not separated because of batch effect. Each color represents a separate batch of samples.

**c.** No difference was observed between male and female mice.

**d,e.** mCherry protein expression perfectly reports *Cyp11a1* mRNA expression. **d.** mCherry protein expression according to the FACS data. **e.** *Cyp11a1* mRNA expression in the scRNA-seq data.

**f.** Expression of cell type specific signature genes that were used to annotate the clusters.

**g.** Marker gene expression in identified clusters.

**h.** Expression level of *Cyp11a1* correlated transcription factors. Pearson correlation was used to identify the *Cyp11a1* correlated genes and pySCENIC was used to identify the potential transcription factors that regulate Cyp11a1 expression.

**i-j.** Differential expression of genes in closely related clusters.

**i.** Pairwise comparison of differential gene expression between three clusters of T helper 2 cells (i.e. Clusters 0, 2 and 3)

**j.** Pairwise comparison of differential gene expression between two clusters of mast cells (Clusters 6 and 9) and basophil/eosinophil cluster (Cluster 7)

k. Expression of glucocorticoid receptor, *Nr3c1*, that is required for glucocorticoid function.

### Extended Data Figure 4

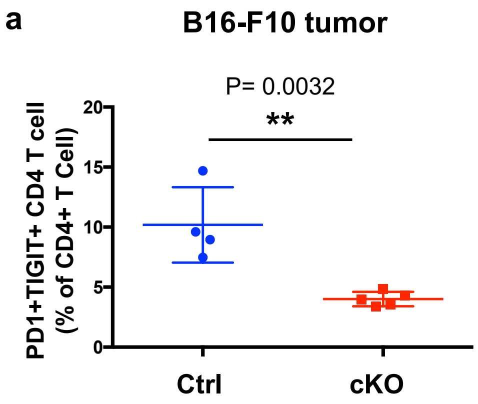


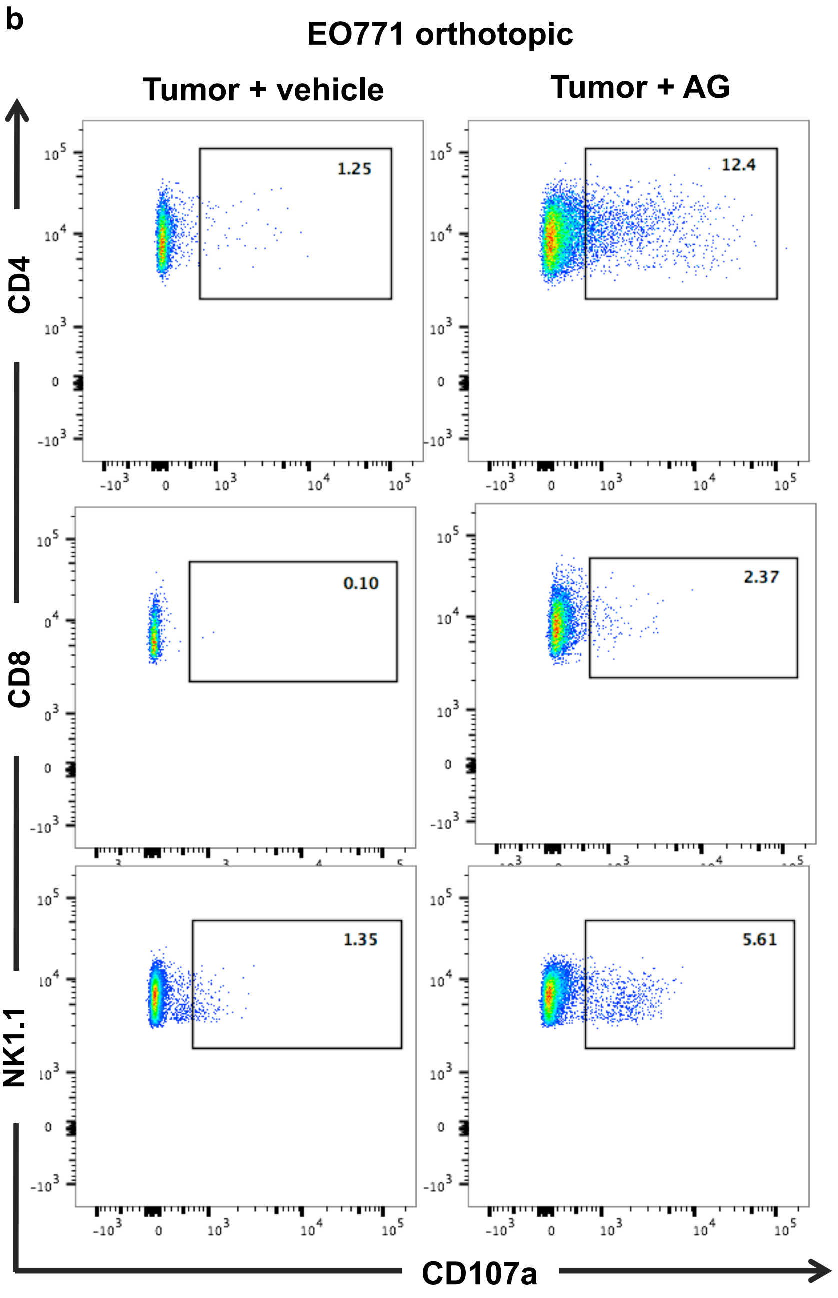

**Extended Data Figure 4. Inhibition of T cell steroidogenesis stimulates anti-tumor immunity**

**a.** Co-inhibitory cell surface receptor PD1 and TIGIT co-expression on tumor infiltrating $CD4^+$ T cells analyzed by flow cytometry after 12 days post B16-F10 inoculation. Gating: All cells > singlets > live cells > CD4+ T cell > PD1, TIGIT.

**b.** Representative FACS profile of Figure 5j, k, l that show degranulation of intratumoral CD4, CD8 T cells and NK cells.
